# Differences in the neural representation of binaural sound localization cues in the auditory midbrain of chicken and barn owl

**DOI:** 10.1101/2023.11.13.566834

**Authors:** Roberta Aralla, Claire Pauley, Christine Köppl

**Affiliations:** Department of Neuroscience, School of Medicine and Health Sciences, Research Center for Neurosensory Sciences and Cluster of Excellence “Hearing4all” Carl von Ossietzky University, 26129 Oldenburg, Germany

## Abstract

The sound localisation behaviour of the nocturnally hunting barn owl and its underlying neural computations is a textbook example of neuroethology. Differences in sound timing and level at the two ears are integrated in a series of well characterised steps, from brainstem to inferior colliculus, resulting in a topographical neural representation of auditory space. It remains an important question of brain evolution how this specialised case derived from a more plesiomorphic pattern. The present study is the first to match physiology and anatomical subregions in the non-owl avian inferior colliculus. Single-unit responses in the chicken inferior colliculus were tested for selectivity to different frequencies and to the binaural difference cues. Their anatomical origin was reconstructed with the help of electrolytic lesions and immunohistochemical identification of different subregions of the inferior colliculus, based on previous characterisations in owl and chicken. In contrast to barn owl, there was no distinct differentiation of responses in the different subregions. We found neural topographies for both binaural cues but no evidence for a coherent representation of auditory space. The results are consistent with previous work in pigeon inferior colliculus and chicken higher-order midbrain and suggest a plesiomorphic condition of multisensory integration in the midbrain that is dominated by lateral panoramic vision.

## Introduction

Of all the sensory systems of vertebrates, the lower auditory ascending pathway is arguably the most complex, with many brainstem nuclei processing different aspects of the input provided by the auditory nerve (e.g., Moss & Carr, 2013; Pickles, 2015). At the level of the *inferior colliculus* (IC), processing streams then converge again. The IC serves as a central hub for auditory processing and an obligatory relay for nearly all ascending auditory information, suggesting complex higher-order processing. A diversity of neural responses are known, and functions have been suggested, which range from simple auditory responses to the representation of auditory space or characteristic features of vocalizations, and to novelty detection and multisensory integration (reviewed by, e.g., Dillingham et al., 2017; Konishi, 2003; Malmierca et al., 2019; Palmer & Kuwada, 2005; Woolley & Portfors, 2013).

In all vertebrates, IC (also known as *nucleus mesencephalicus lateralis pars dorsalis* – MLd – in birds, and as *torus semicircularis* – ToS – more generally in non-mammalian vertebrates) comprises two or more subdivisions that can be identified by anatomical and physiological criteria (reviewed by, e.g., Bass et al., 2005; Belekhova et al., 2020; Covey & Carr, 2005; Ehret & Schreiner, 2005; Endepols et al., 2000; Grothe et al., 2004). Typically, a principal distinction is made between a core region (often termed the central nucleus) that receives the bulk of ascending brainstem projections, and a surrounding belt (often subdivided in to several regions termed external or cortical) of higher order. Beyond that, however, there is no consensus across species on the number, nomenclature, and definition of subregions.

The core and belt regions of IC represent different hierarchies and are believed to have different roles in sensory processing. A common pattern found in both mammals and birds is the purely auditory nature of the central nucleus, and the selectivity of its neurones for binaural difference cues. The central nucleus always receives direct inputs from the auditory brainstem and has neurones that are narrowly frequency tuned and tonotopically organized along the dorso-ventral axis (reviewed by, e.g., Casseday et al., 2002; Ehret & Schreiner, 2005; Ito et al., 2019; Palmer & Kuwada, 2005; Singheiser et al., 2012; Yin & May, 2005). Less is known about the IC’s cortical regions. In mammals, these regions typically receive inputs from the central nucleus and other, non-auditory sensory areas, as well as prominent descending projections from the auditory cortex. They are believed to be involved in higher-order processes such as multimodal integration or sensory novelty detection (reviewed by, e.g., Ito & Malmierca, 2018; Winer & Schreiner, 2005).

Studies of the IC of birds has a long history. The arguably most influential work was carried out on the processing of binaural cues for sound localization in the barn owl (for a historical review see Carr & Pena, 2016). The barn owl is an auditory specialist that has hypertrophied auditory nuclei when compared to other birds (Gutiérrez-Ibánez et al., 2011; Kubke et al., 2004), and the nuclei are therefore easier to identify and target. In the barn owl, IC subregions can be readily distinguished by anatomical features, as well as through the different selectivities of their neurones for frequency and for sound-localization cues. In the barn owl IC, the separate brainstem processing streams for the two principal binaural cues, interaural time (ITD) and interaural level (ILD) difference, terminate and subsequently converge for the first time, ultimately generating a two-dimensional neural representation of auditory space (reviewed by, e.g., Kettler & Carr, 2019; Konishi, 2003; Singheiser et al., 2012).

We use here a nomenclature for the IC subregions that was introduced for the barn owl and subsequently adopted for the chicken. It identified a central (ICC), a superficial (ICS), and an external (ICX) nucleus (Knudsen, 1983; Niederleitner & Luksch, 2012; Wagner et al., 2003; but see Logerot et al., 2011; Puelles et al., 1994), for different definitions). In the barn owl, further subdivisions of ICC were defined, based on separate lower brainstem projections to a core (ICCc) and a shell (ICCs). The shell was further differentiated into lateral (ICCls) and medial parts (ICCms; Adolphs, 1993a; Takahashi & Konishi, 1988; Takahashi et al., 1989).

The most striking characteristic of the barn owl IC is its map of auditory space in the ICX (Knudsen and Konishi, 1978). Here, “space-specific” neurones (a term that was introduced later; Takahashi et al., 1984) are excited exclusively by sound coming from a well circumscribed region of auditory space, i.e., individual neurones are selective for a restricted range of space in both azimuth and elevation. They are topographically arranged in a two-dimensional space map: azimuthal (horizontal) space is represented along the rostro-caudal axis, elevational (vertical) space along the dorso-ventral axis (Knudsen & Konishi, 1978). In the space map of barn owl, sensitivity to ITD and ILD has been shown to underlie the selectivity for azimuth and elevation, respectively (Takahashi et al., 1984). To date, a similar two-dimensional map has not been found in any other species. Simpler spatial representations, in which IC neurones with spatially restricted auditory receptive fields were arranged into one-dimensional maps of azimuth have been found in several other owl species (Volman & Konishi, 1990), in another bird of prey (Calford et al., 1985), and in the pigeon (Lewald, 1990).

Overall, the prominent example of the barn owl has led to a distorted perception that often portrays this clearly fascinating but specialised bird as representative for the processing of auditory spatial information in birds. In this study, we investigated the anatomy and physiology of IC subregions in a generalist bird, the chicken (*Gallus gallus*). The anatomy of the chicken IC, as well as the neural responses along the auditory pathway including the IC, share salient similarities with the well-known specialized system in the barn owl. The same IC subdivisions can be identified anatomically in chicken and barn owl (Niederleitner & Luksch, 2012). Moreover, in the chicken brainstem, the presence of delay lines and topographic maps of ITD suggest a Jeffress-like coding model, as demonstrated first in barn owl and subsequently in other birds (Ashida & Carr, 2011). Chicken IC neurones predominantly respond to sounds from contralateral auditory space, show a clear dorso-ventral tonotopy, and most neurones are selective for ITD, ILD or both (Aralla et al., 2020). However, the anatomical orientation of the chicken’s ITD maps in the brainstem are different to those in the barn owl (Köppl & Carr, 2008). In these respects, the chicken rather resembles less specialized species of birds (MacLeod et al., 2006) and other archosaurs (Kettler & Carr, 2019). Importantly, for the coding of sound location in the chicken IC, our single-unit data suggested complementary roles for ITD and ILD across frequencies (Aralla et al., 2020). This is in line with the duplex theory (Rayleigh, 1907), and was also postulated for other generalist birds such as pigeons (Lewald, 1988, 1990), and for mammals (e.g., Keating et al., 2014). These characteristics made us choose the chicken as a suitable animal model for studying plesiomorphic mechanisms in the auditory system, and for deriving a default layout of the avian auditory system.

The present study aims to bring together the anatomy and the physiology of IC in this generalist bird. To this end, we defined, in three-dimensional space, the histological subdivisions of the chicken IC, differentiating between the core (ICCc) and the shell (ICCs) of the ICC, as well as ICX. Furthermore, we reconstructed individual electrode tracks from previous electrophysiological experiments in 3-dimensional space and assigned the recorded auditory responses to specific IC subregions. This allowed us to correlate the subregions’ selectivity for frequency and for binaural cues.

## Results

The chicken IC is located in the midbrain, surrounded by the optic tectum. In particular, it lies immediately under the lateral recess of the aqueduct (Iraq; Puelles et al., 2019), bordered dorsally and laterally by the intercollicular area and ventrally by the pre- isthmic nuclei. We present the results of a histological analysis performed on the left-side IC of seven chicken midbrains. All brains were Nissl-stained, which generally showed the electrolytic lesions placed during previous recording sessions most clearly. In five of seven brains, alternating tissue sections were antibody-labelled (see Methods), to enable the identification of IC subregions. Our samples were routinely labelled with a pair of primary antibodies that targeted different structures in a complementary way: anti-CKII and anti-PV most strongly labelled ICX or ICCc, respectively (Table 1). Early attempts with anti-calretinin showed only relatively weak labelling of ICCc and this was not further pursued. In the final sample reported here, one anti-calretinin-only labelled brain, as well as two Nissl-only stained brains were included (see Methods for details), since we had valuable physiology from these individuals.

**Table 1:** Primary antibodies and their labelling of IC subregions.

|  |  | ICCc | ICCs | ICX |
| --- | --- | --- | --- | --- |
| anti-PV | cell bodies | very dark | dark | unstained |
|  | neuropil | very dark | dark | unstained |
| anti-CKII | cell bodies | unstained | dark | very dark |
|  | neuropil | unstained | light | very dark |

### Anatomical characterization of IC subregions

The core of ICC was located in the centro-ventral region of IC (Fig. 1B, 2B). Medially, laterally, and dorsally it was always surrounded by the shell (ICCs). ICCc was strongly labelled by anti-PV, where both cell bodies and neuropil appeared black (Fig. 2C), presenting a gradient between ICCc and ICCs. Anti-CKII left the ICCc only lightly labelled or completely unlabelled (Fig. 1).

**Figure 1:**
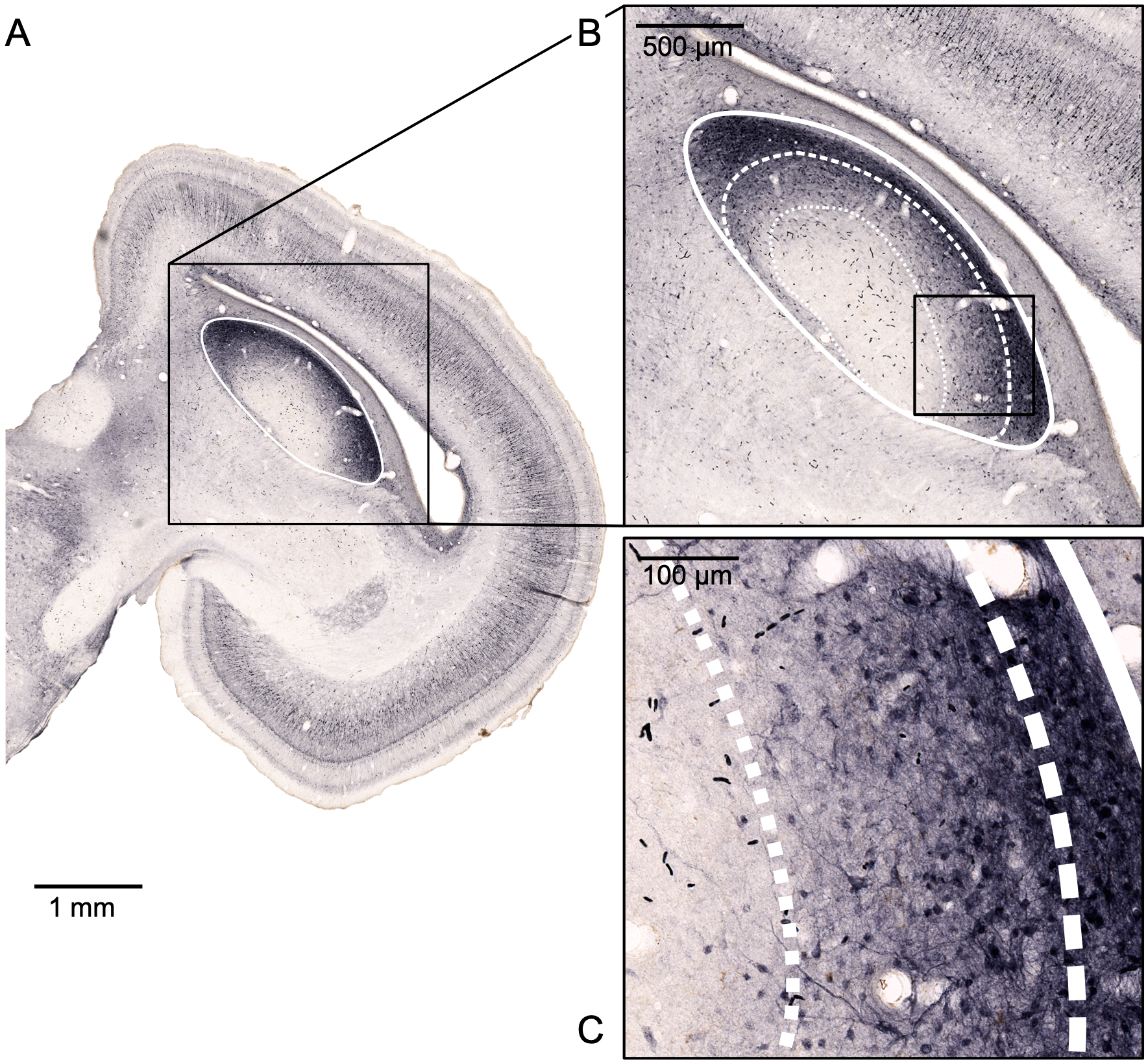
Differential immunolabelling of IC subregions with anti-CKII. (appearing in black). Panel A shows a histological cross-section of the midbrain at a position 1530 µm (84%) rostral of IC’s caudal outer border. The solid white line indicates the outline of IC. The area indicated by the black box is shown enlarged in panel B, with the outlines of the three subregions ICCc, ICCs and ICX as dotted, dashed and continuous white lines, respectively. Panel C shows the area indicated by the black box in B at higher magnification. Note the differential CKII-labelling providing a clear contrast between the darkly labelled ICX and the unlabelled ICCs, and a gradient of labelling intensity of somata and neuropil in the ICCs.

**Figure 2:**
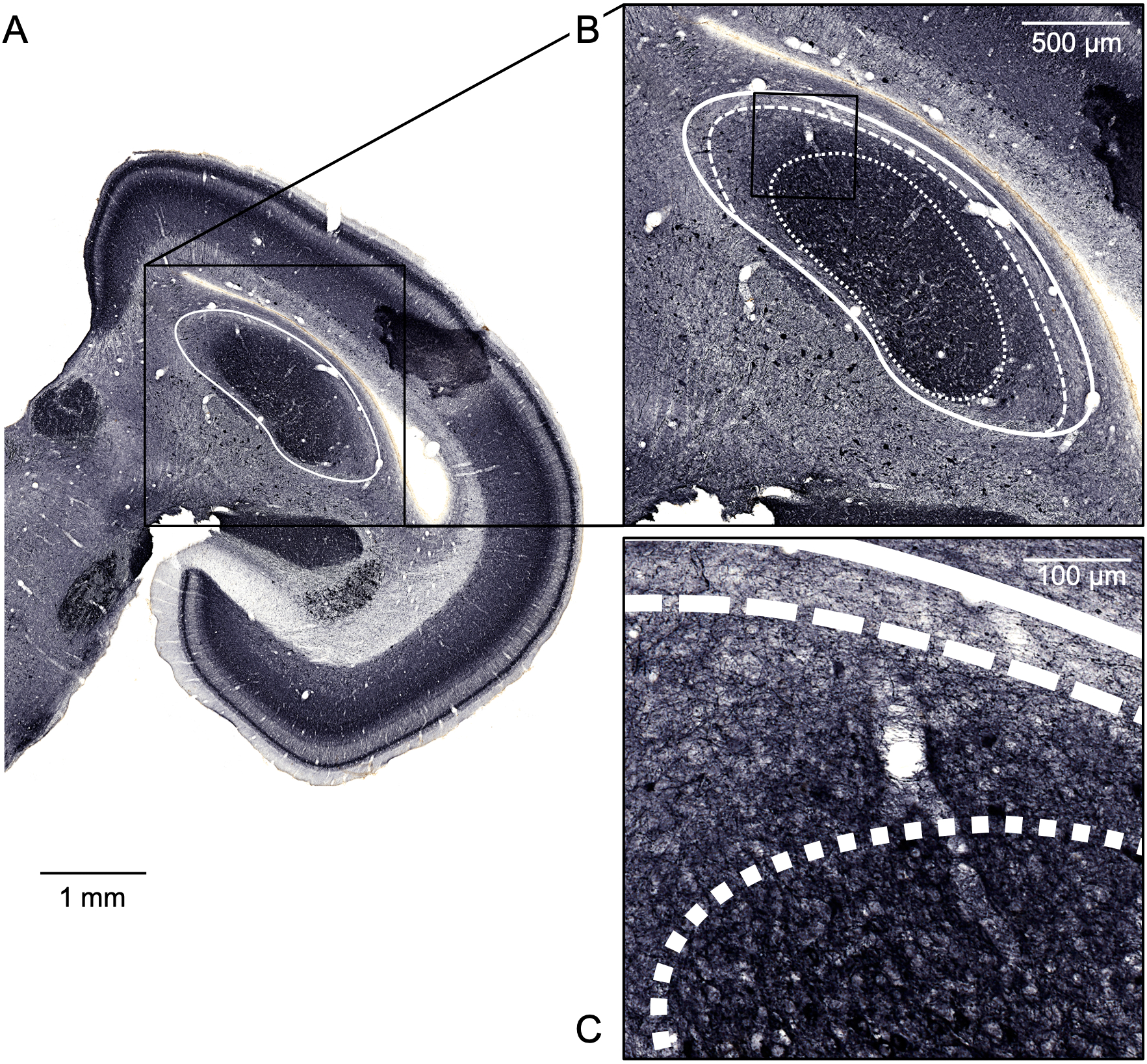
Differential immunolabelling of IC subregions with anti-parvalbumine. (appearing in black). Panel A shows a histological cross-section of the midbrain at a position 1470 µm (80%) rostral of IC’s caudal outer border. The solid white line indicates the outline of IC. The area indicated by the black box is shown enlarged in panel B, with the outlines of the three subregions ICCc, ICCs and ICX as dotted, dashed and continuous white lines, respectively. Panel C shows the area indicated by the black box in B at higher magnification. Note that the borders of the three subregions are defined differentially by parvalbumine labelling, which shows a complementary pattern to the CKII-label (Fig. 1). Between the unlabelled ICX and the darkly labelled ICCc there is a progressive darkening of the label of somata and neuropil in the ICCs.

The shell of ICC enclosed the core almost completely and also defined the ventral border of the IC. The shell was usually found along the entire caudo-rostral extent of the IC. In ICCs, anti-CKII and anti-PV darkly labelled only the somata (Fig. 1, 2). Interestingly, the labelling of the shell typically showed an increasing (with anti-CKII, Fig. 1B) or decreasing (with anti-PV, Fig. 2B) gradient of intensity, from the core outwards. In our samples, it was not possible to make any morphological distinction between lateral and medial shell. The border between ICCs and ICX was precisely defined by anti-PV and anti-CKII. Labelled somata were typically round or oval shaped (Fig. 1C, 2C), and no dendritic fields were detectable.

For most of the caudo-rostral extent of IC, ICX was a thin layer forming the outer cover of the IC on its medial, dorsal, and lateral sides. ICX borders were delineated by anti-PV and anti-CKII; while anti-CKII labelled both cell bodies and neuropil of the ICX very darkly (Fig. 1), anti-PV left ICX unlabelled (Fig. 2). Thus, these two antibodies clearly marked the ventral border between ICCs and ICX, and between ICX and the ICS or the parenchymal tissue dorsally. Cell bodies labelled in the ICX appeared very dense and compact, and had a round or oval shape (Fig. 1C).

### Linear dimensions and volumetric size of IC subregions

This study performed for the first time a quantitative analysis of the chicken auditory midbrain, aiming to characterize the typical dimensions of its subregions. In the immunolabelled samples, the linear extents of all of IC’s subregions were measured along the three axes – medio-lateral, ventro-dorsal and caudo-rostral – while in the remaining two brains only the outer borders of IC could be defined. Across all seven animals, IC had a median total caudo-rostral extent of 1830 µm (quartiles 25%=1395 µm; 75%=2145 µm), a median total medio-lateral extent of 3321 µm (quartiles 25%=3199 µm; 75%=3448 µm) and a median total ventro-dorsal extent of 2239 µm (quartiles 25%=1767 µm; 75%=2614 µm). A summary of all subregions’ measurements is provided in Table 2; Fig. 3A displays the data graphically.

**Figure 3:**
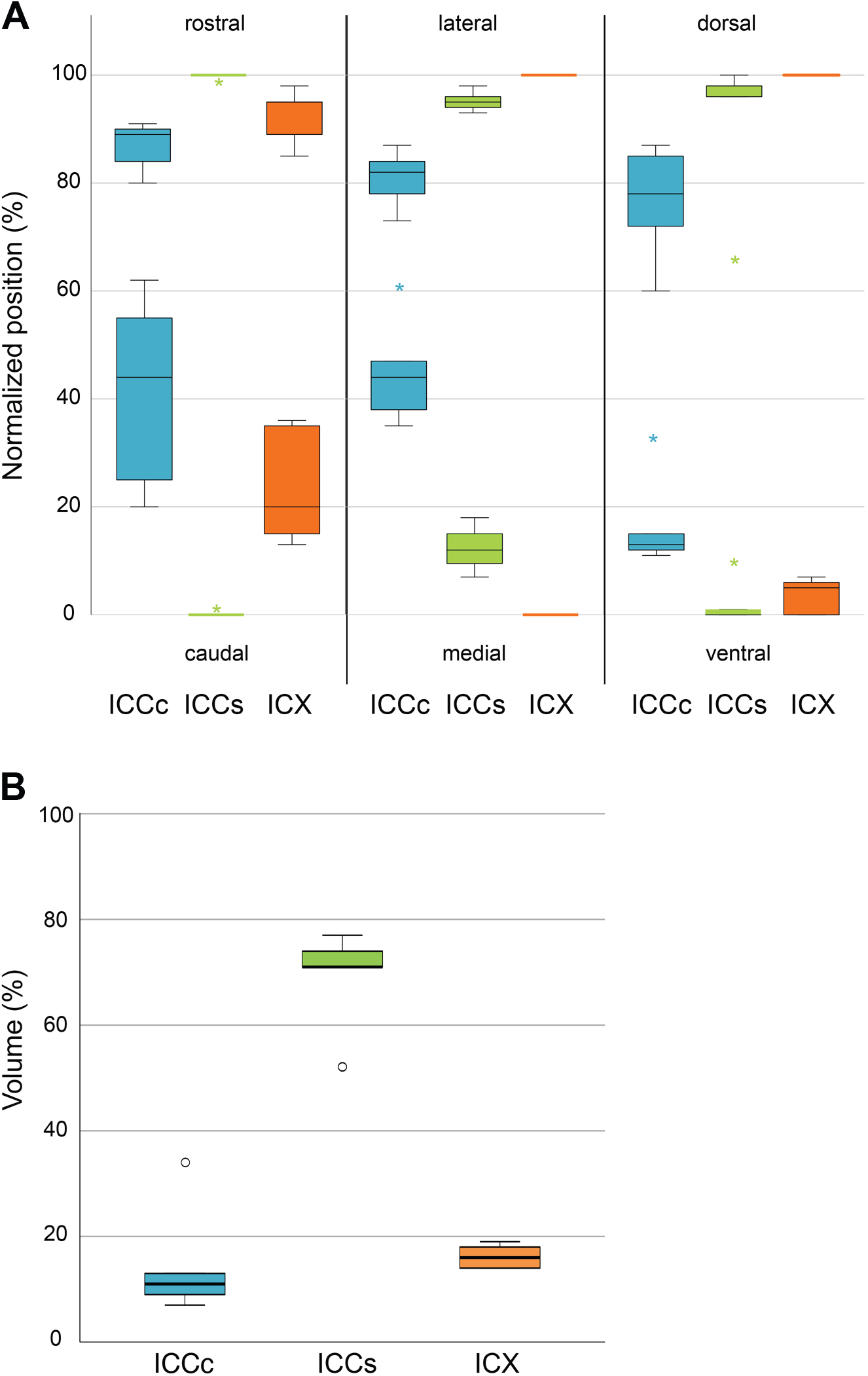
Relative sizes and volumes of IC subregions. Data for the ICCc are shown in blue, for ICCs in green, and for ICX in orange. A: Outer borders of the three quantified subregions of IC. Boxplots represent the outer limits of the three subregions along the caudo-rostral, medio-lateral and ventro-dorsal axis, respectively, normalized to the total extent of IC along each axis. Lower boxplots indicate caudal, medial and ventral limits, while upper ones indicate rostral, lateral and dorsal ones. Each box represents the interquartile range, with the median indicated by the horizontal line, whiskers represent the data range (n = 5 brains). The color-coded asterisks indicate the outliers. B: Boxplot of normalized volumes of the three IC subregions, shown as percentages of the total IC volume. Note that the ICCs is by far the largest subregion.

**Table 2:** Quantitative analysis of IC’s subregions along the three dimensions. Each distance is given relative to the caudal, medial or ventral outer border, respectively, of the IC. Bold numbers indicate the limits of the subregions that also define the outer borders of IC. Medians, as well as 25^th^ and 75^th^ percentile values are listed.

|  |  |  | ICCc |  | ICCs |  | ICX |  |
| --- | --- | --- | --- | --- | --- | --- | --- | --- |
|  |  |  | µm | % | µm | % | µm | % |
| caudo-rostral axis | Caudal border | median | 870 | 44 | 0 | 0 | 360 | 20 |
|  |  | 25%, 75% | 450; 1260 | 25; 55 | 0 | 0 | 300; 690 | 15; 35 |
|  | Rostral border | median | 1800 | 89 | 1980 | 100 | 1950 | 95 |
|  |  | 25%, 75% | 1650; 1950 | 84; 90 | 1830; 2310 | 100 | 1740; 2160 | 89; 95 |
|  | CR length | median | 750 | 47 | 1980 | 100 | 1290 | 71 |
|  |  | 25%, 75% | 690; 930 | 30; 61 | 1830; 2310 | 100 | 1260; 1380 | 64; 75 |
| medio-lateral axis | Medial border | median | 1394 | 44 | 421 | 12 | 0 | 0 |
|  |  | 25%, 75% | 1288; 1551 | 38; 47 | 343; 564 | 12; 18 | 0 | 0 |
|  | Lateral border | median | 2489 | 82 | 2835 | 95 | 2957 | 100 |
|  |  | 25%, 75% | 2427; 3034 | 78; 84 | 2789; 3201 | 94; 96 | 2956; 3395 | 100 |
|  | ML length | median | 655 | 16 | 1271 | 43 | 2246 | 76 |
|  |  | 25%, 75% | 462; 907 | 14; 30 | 400; 2149 | 13; 68 | 1395; 2718 | 43; 92 |
| ventro-dorsal axis | Ventral border | median | 265 | 13 | 0 | 0 | 92 | 5 |
|  |  | **25%, 75%** | 239; 438 | 12; 15 | **0; 30** | **0; 1** | 0; 100 | 0; 6 |
|  | Dorsal border | **median** | 1540 | 78 | 1784 | 98 | **1798** | **100** |
|  |  | **25%, 75%** | 1404; 2106 | 72; 85 | 1726; 2303 | 96; 98 | **1784; 2400** | **100** |
|  | VD length | **median** | 694 | 32 | 1012 | 38 | 1549 | 67 |
|  |  | **25%, 75%** | 515; 1014 | 23; 45 | 294; 1572 | 14; 83 | 582; 1621 | 30; 87 |

As described above, ICCc formed the inner-most part of the IC and did not touch the outer borders of the IC on any axis. ICCc’s relative extent typically comprised 45% of the total caudo-rostral extent of the IC (between 44-89% from the outer caudal border), and 65% of its total ventro-dorsal extent, but only 38% of its medio-lateral extent. ICCs and ICX wrapped around the core like the layers of an onion, albeit not uniformly. A thick layer of shell typically bordered the core, and was itself enclosed by a thinner layer of ICX. The outer-most borders of the IC on the medial, lateral and dorsal extremes were defined by ICX, but on the caudal, rostral and ventral extremes by ICCs (Table 2 and Fig. 3A).

The above qualitative and quantitative analyses of the chicken IC provided the necessary information for a three-dimensional reconstruction of IC’s subregions and derivation of their volumes. The entire IC had a median volume of 2.39 mm³ (n=5; quartiles 25%= 2.28 mm³; 75%= 2.69 mm³; range 1.87 mm³ to 2.80 mm³). Of the three subregions, the ICCc showed the smallest volume (Fig. 3B), occupying only 11% of the entire IC (median = 0.24 mm³; 25th quartiles = 0.23 mm3 and 9%; 75th quartiles = 0.3 mm³ and 13%). The core was followed in size by ICX, which took up 16% of the entire IC (median = 0.38 mm³; 25th quartiles = 0.34 mm³ and 14%; 75th quartiles = 0.38 mm³ and 18%). The largest subregion by far was the ICCs, which comprised 71% of the entire IC (median = 1.63 mm³; 25th quartiles = 1.52 mm³ and 71%; 75th quartiles = 2.06 mm³ and 74%).

### Physiological characterization of single units in IC

During the electrophysiological experiments under dichotic stimulation, each single-unit recording was characterized by its best frequency (BF) and sensitivity to differences in interaural time (ITD) and level (ILD) (Aralla et al., 2020). Briefly, frequency responses were classified as low (LF, BF ≤ 2 kHz), high (HF, BF > 2 kHz), or broadly tuned/not frequency selective (broadband, BB). ITD selectivity, found in 65% of units, was typically phase-ambiguous. As it is controversial whether the neurones’ maximal discharge or the point of maximal change in discharge rate is the relevant parameter that encodes sound location, Aralla et al. (2020) determined both but could not clearly identify the more physiologically relevant one. Here, we will only refer to best ITD (corresponding to the point of maximal discharge, closest to zero ITD). Note that the likely physiological ITD range, i.e. the maximal ITDs experienced by an adult chicken with natural free-field stimulation, is between ±130 µs and potentially more than ±200 µs at frequencies <1kHz (Köppl, 2019; Schnyder et al., 2014). For ILD, a surprising variety of selective response types was found (in total, 76% of all units), some of which could be plausibly explained by the purely physical interactions between the internally coupled middle ears of chickens and were thus classified as monaural (response type “dip”). The remaining ILD response types (“contra-dominated”, “ipsi-dominated”, “V-shaped” and “open-bell-shaped”) likely included synaptic binaural integration. We defined a best ILD for these which, depending on the response type, was either the point of maximal or minimal discharge, or the point of maximal slope. To put this parameter into a natural localization context, an adult chicken head was shown to cause free-field ILDs of ±8 dB via sound shadowing (Schnyder et al., 2014); this would be further enhanced by the internally coupled middle ears (Larsen et al., 2016), but precise numbers have not been determined for the chicken.

### 3D-reconstruction of electrode tracks

Here, we now add the anatomical position within the IC of each recording site, carefully reconstructed in each individual brain, and thus correlate their physiological features with IC’s subregions. The histological analysis recovered 24 (of 28 placed) electrolytic lesions (examples in Fig 4A-C). Together, the recovered lesions enabled the reconstruction of 14 electrode tracks, with a total of 114 single-unit recordings, which represent the majority of the data reported by Aralla et al. (2020; 139 units). Along the medio-lateral axis of IC, the recording sites were located between 32% and 86% from the medial border, along the ventro-dorsal axis between 16% and 94% from the ventral border, and along the caudo-rostral axis, between 28% and 93% from the caudal border. Thus, the IC was sampled quite comprehensively, but not exhaustively. Furthermore, within these outer limits, the distribution of recording sites was clearly biased towards the lateral half of the IC (see also later Figs. 7 and 9). Overall, this indicates that the ICX and the medial part of ICCs are underrepresented.

**Figure 4:**
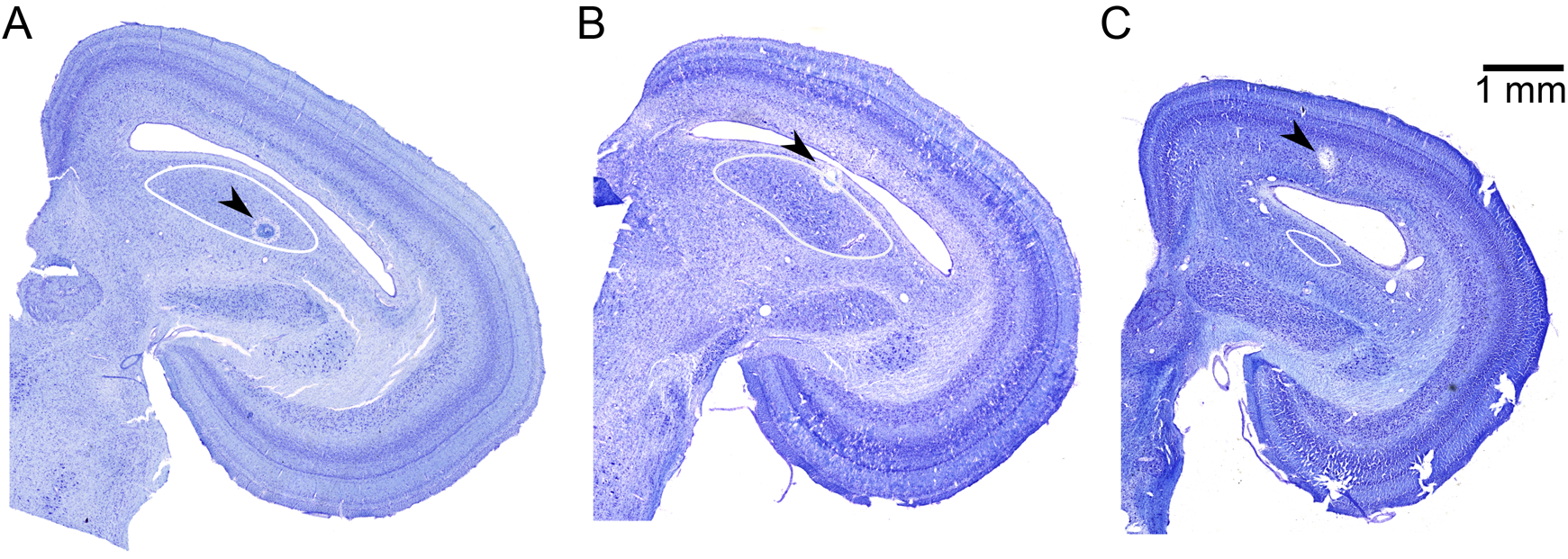
Examples of electrolytic lesions. Three Nissl-stained histological sections illustrating different positions of lesions relative to IC. Lesions, indicated by the black arrowheads, were either found inside (A), on the border (B) or outside (C) the IC (outlined by the solid white line). The sections are shown uniformly oriented, with the brains’ midlines vertical, but represent different positions along the caudo-rostral axis of three different midbrains: A: 2490 µm (85%), B: 1830 µm (79%), and C: 2430 µm (100%) rostral of IC’s caudal outer border.

Note that we distinguished between unambiguously and plausibly located recording sites, and also carefully considered alternative subregion allocations for recording sites that fell close to subregion borders (see Methods). Between 25 and 28 sites fell within the ICCc (20 of those unambiguously), between 66 and 71 fell within the ICCs (41 unambiguously) and five to eight within the ICX (none unambiguously). Between 10 and 15 recording sites were located just outside the borders of IC, at up to 630 µm distance. Below, the distribution of the previously identified response types (Aralla et al., 2020) across the IC subregions will be examined. None of the principal conclusions was affected by the ambiguities in locating the recording sites. Therefore, the numbers given below refer to the pooled sample (unambiguously and plausibly located) and the primary allocated subregion of each recording site, unless explicitly stated otherwise.

### Distribution of unit response characteristics across the different IC subregions

Units in ICCc were all frequency-selective, with an overrepresentation of LF units (22 of 28; 79% vs. 69% in the entire IC sample; Fig.5). Furthermore, most units in ICCc were ITD-sensitive (19 of 26 tested), consistent with its low-frequency bias (Aralla et al., 2020). The best ITD of LF neurones ranged between -200 (ipsilateral leading) and 917 µs (contralateral leading), while the range of best ITDs was much smaller for the three HF neurones, between 99 and 139 µs. ILD sensitivity was found at 19 of 24 recording sites tested in ICCc, which was a somewhat lower proportion (79%) than overall (87%) and, again, consistent with ICCc’s low-frequency bias (Aralla et al., 2020). ILD responses were classified as “contra-dominated” for all HF-tuned units located in ICCc (6 of 19), while LF-units showed more heterogeneous ILD responses, of the “v-shaped” (8), “ipsi-dominated” (1) or “dip” (4) types. Over all response types, best ILD ranged from -11 to 6 dB, and almost identically for LF and HF units. Fourteen ICCc units were sensitive to both ITD and ILD, which was a typical proportion for the entire IC.

**Figure 5:**
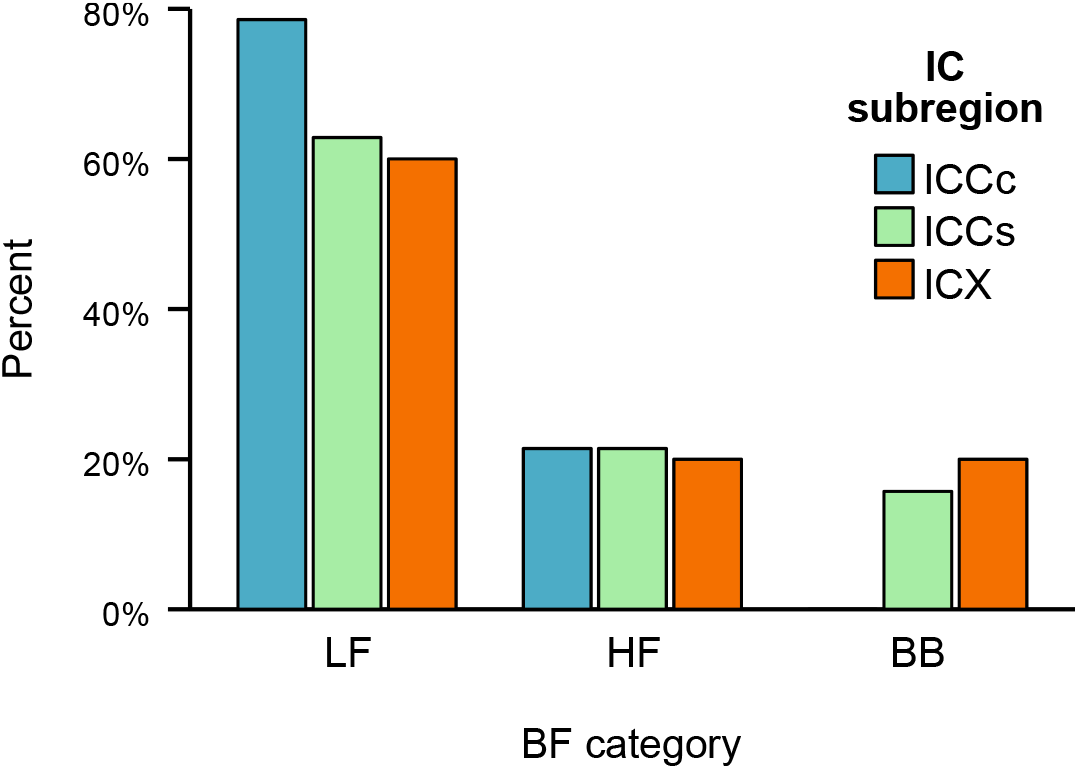
Distribution of units with different frequency selectivity across the IC subregions. Units were categorized into low-frequency (LF, best frequencies ≤ 2 kHz), high-frequency (HF, best frequency > 2 kHz) and broadband (BB, i.e. not frequency selective). Data for the three IC subregions are colour coded: ICCc in blue (n = 28), ICCs in green (n = 70) and ICX in orange (n = 5). Note that bar height represents the percentage within each subregion, which emphasises differences in the distribution of response classes across subregions.

Recording sites located in the shell of the ICC showed LF selectivity in 44 of 71 cases, while 15 were HF selective, and 11 were classified as broadband (BB); one unit was not tested for BF. This amounts to a slight underrepresentation of LF units and a clear overrepresentation of BB responses relative to the entire IC sample (Fig. 5). Forty of 60 ICCs units proved to be selective for ITD sensitivity. Their best ITD ranged broadly for LF units between 2 and 771 µs, for HF units between 20 and 135 µs, and for BB units between 0 and 362 µs. Selectivity for ILD was very common in ICCs and slightly overrepresented (92% of units tested vs. 87% overall), consistent with the relative predominance of HF and BB units (Aralla et al., 2020). The ILD responses in ICCs were mostly of the “contra-dominated” type (40 of 55 tested; Fig. 6). ILD responses of the “ipsi-dominated” type were found in four cases, “v-shaped” in only one case, and “dip” in six cases, the latter all tuned to LF. All four ILD responses of the overall rarest “open-bell-shaped” type were located in the ICC shell. However, only one of these could be allocated unambiguously, while three of them alternatively fell outside of IC. “Open-bell-shaped” ILD responses were thus always found near the outer edge of IC. Thirty-five ICCs units were sensitive to both ITD and ILD, which was an overall typical proportion.

**Figure 6:**
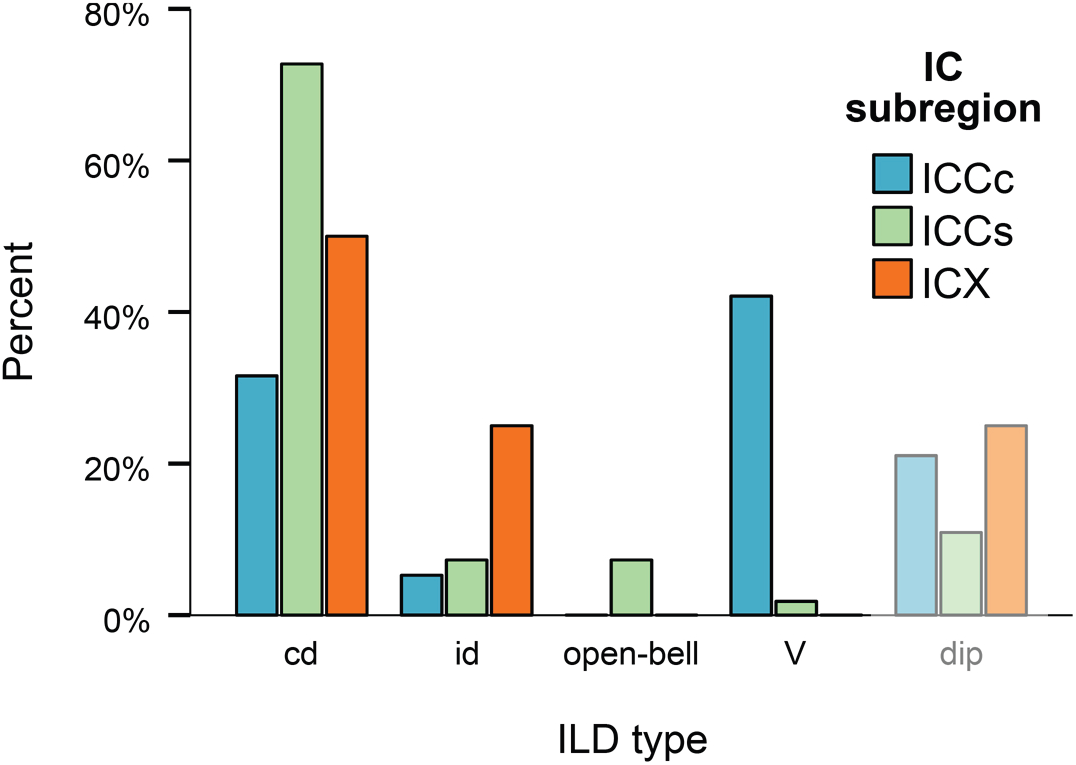
**Distribution of ILD-sensitive units across the IC subregions**, separated according to ILD response type: “contra-dominated” (cd), “ipsi-dominated” (id), “open-bell-shaped”, “v-shaped” and “dip” (greyed out to indicate that these are likely monaural responses, arising from physical interaural interaction; (Aralla et al., 2020). Data for the three IC subregions are colour coded: ICCc in blue (n = 19), ICCs in green (n = 55), and ICX in orange (n = 4). Note that bar height represents the percentage within each subregion, which emphasises differences in the distribution of response classes across subregions.

The five recording sites plausibly located in the ICX showed all types of frequency selectivity: three LF, one HF, and one BB (Fig. 5). Considering the small sample size, it seems noteworthy that one of the fairly rare BB units was among them. Three ICX neurones were selective for both ITD and ILD, including the BB unit, and one each was sensitive to only ITD or ILD. Several types of ILD responses were located in ICX: two “contra-dominated“, and one each of the “ipsi-dominated” and “dip” types (Fig. 6).

In summary, all subregions represented a wide tonotopic range of frequency-selective units. However, broadly tuned or frequency-unselective units were only found in ICCs and ICX. ICCc neurones, in addition to always being narrowly frequency selective, showed an overrepresentation of LF sensitivity. In terms of ITD-selectivity, ICCc and ICCs showed slight biases that were complementary and also correlated with the higher likelihood for ITD-selectivity in LF-units (Aralla et al., 2020): ICCc had a higher proportion of ITD-selective neurones than ICCs. The reverse was true for ILD-selective units, where ICCc had a lower proportion than ICCs, again consistent with their BF biases (Aralla et al., 2020). The most striking differentiation was observed in the distribution of ILD response types (Fig. 6). “Contra-dominated” and “ipsi-dominated” ILD responses were overrepresented in the shell of ICC, and all “open-bell-shaped” responses were located in the shell. In contrast, “V-shaped” ILD responses were nearly exclusively located in the core of ICC. The small ICX sample showed a surprising variety of physiological responses and no clear over- or under-representations of specific types could be identified.

### Topographic representations of frequency and binaural cues

Correlations with the location of neurones along all three anatomical axes were explored for BF, best ITD, and best ILD, separately in ICCc and ICCs. Our ICX sample was too small for a meaningful statistical analysis and is only displayed graphically in Figs 7 and 8.

**Figure 7:**
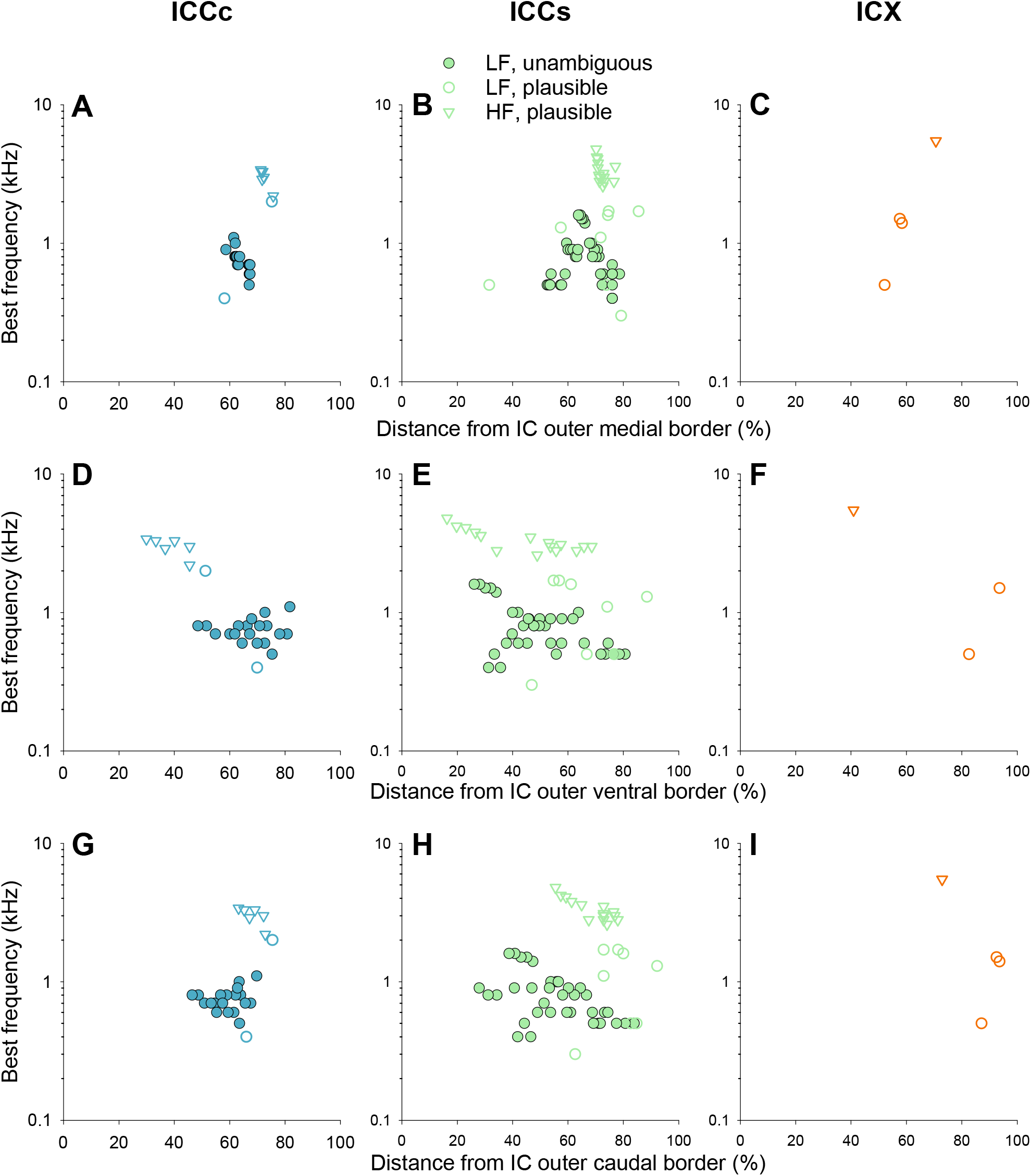
Tonotopic gradients in IC. The individual panels show the BFs of frequency-selective neurones, as a function of their anatomical positions along the three axes, medio-lateral (A-C), ventro-dorsal (D-F) and caudo-rostral (G-I), and separated according to the three IC subregions, ICCc (A, D, G), ICCs (B, E, H) and ICX (C, F, I). Filled symbols indicate recording sites classified as “unambiguous”, while open symbols indicate “plausible” (see Methods). Circles show data for LF units, triangles for HF units. Different colours distinguish the IC subregions, according to the uniform scheme used in all figures. The data supported a clear tonotopic topography along the ventro-dorsal axis (D, E) in ICCc and ICCs, with additional evidence for a caudo-rostral tonotopic gradient (G, H; details see main text).

**Figure 8:**
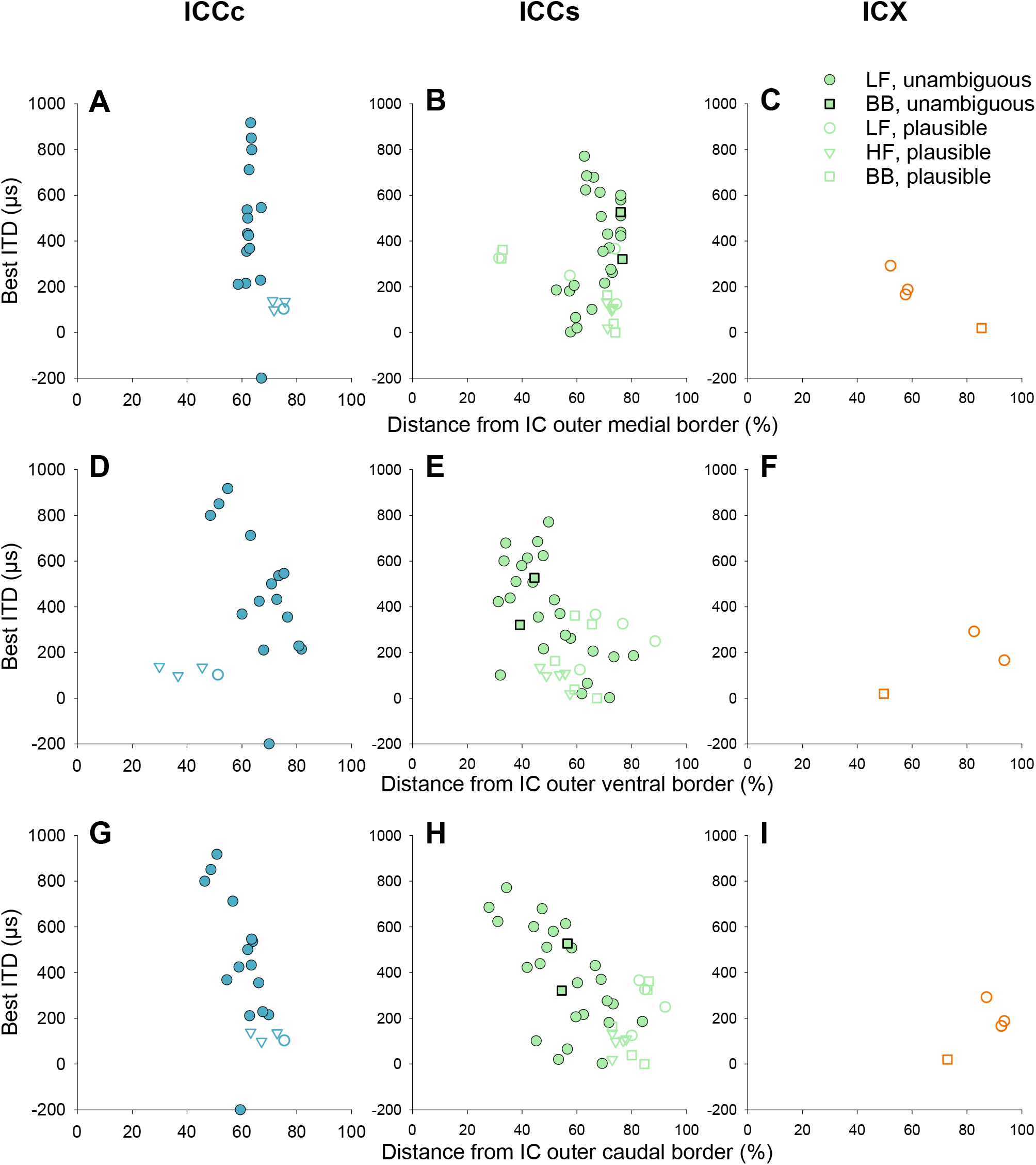
Topographic representation of ITD in IC. The individual panels show the best ITD of individual neurones as a function of their anatomical positions along the three axes, medio-lateral (A-C), ventro-dorsal (D-F) and caudo-rostral (G-I), and separated according to the three IC subregions, ICCc (A, D, G), ICCs (B, E, H) and ICX (C, F, I). Filled symbols indicate recording sites classified as “unambiguous”, while open symbols indicate “plausible” (see Methods). Circles show data for LF units, triangles for HF units, and squares for BB (not frequency selective) units. Different colours distinguish the IC subregions, according to the uniform scheme used in all figures. The data supported clear ITD mapping along the ventro-dorsal (D, E) and caudo-rostral axes (G, H), for both ICCc and ICCs (details see main text).

In both ICC subregions, frequency-selective neurones were clearly arranged tonotopically along the ventro-dorsal axis, with BF increasing towards ventral (Fig. 7D, E; Spearman rank correlation between location and ln BF, recording sites classified as “unambiguous” and “plausible” pooled, for ICCc: n = 28, rho = -0.55, p = 0.003, for ICCs: n = 59, rho = -0.32, p = 0.014). Tonotopic gradients along the caudo-rostral or medio-lateral axes were less convincing. Here, the correlations varied with the precise sample (unambiguously and plausibly located, and also alternative subregion allocations). Intriguingly, both ICCc and ICCs showed two clusters of units that were not related to the subregion allocation, separating instead approximately according to BF (Fig. 7D, E). This suggests an extensive spatial overlap of LF and HF units, with tonotopic gradients running parallel to each other (instead of one joint tonotopic gradient) along the ventro-dorsal (Fig. 7D-F) and caudo-rostral axes (Fig. 7G-I). However, the few individual electrode tracks that spanned larger BF ranges showed no evidence for alternating between LF and HF units. Thus, an alternative and more plausible explanation is that, while a typical tonotopic gradient from rostro-dorsal (LF) to caudo-ventral (HF) is shared, individual chicken’s ICC differed in the relative locations of specific BFs along that axis. Such differences could reflect biological variation and/or the different reliabilities of histological reconstruction.

A consistent topographic organization of best ITD was observed along the caudo-rostral axis. Best ITDs increased in both ICCc and ICCs, from values near zero (frontal auditory space) rostrally to contra-leading ear caudally (Fig. 8G, H; Spearman rank correlation, for ICCc: n = 19, rho = 0.64, p = 0.003, for ICCs: n = 40, rho = 0.54, p < 0.001). In ICCs, there was, in addition, an equally robust topography along the ventro-dorsal dimension, with best ITD increasing towards ventral (Fig. 8E; rho = -0.54, p < 0.001). The ITD topographies were clearly dominated by LF units that contributed the majority of ITD-sensitive sites, and all unambiguously located units in ICCc. If only LF units were considered, the ventro-dorsal axis of ICCc also showed a systematic change of best ITD (Fig. 8D; n = 15, rho = -0.56, p = 0.03, unambiguous only). These population trends were also consistent with the progression of best ITD along individual electrode tracks (Fig. 9), but note that the sequence was never completely unidirectional (examples in Fig. 9B, E, H, K).

**Figure 9:**
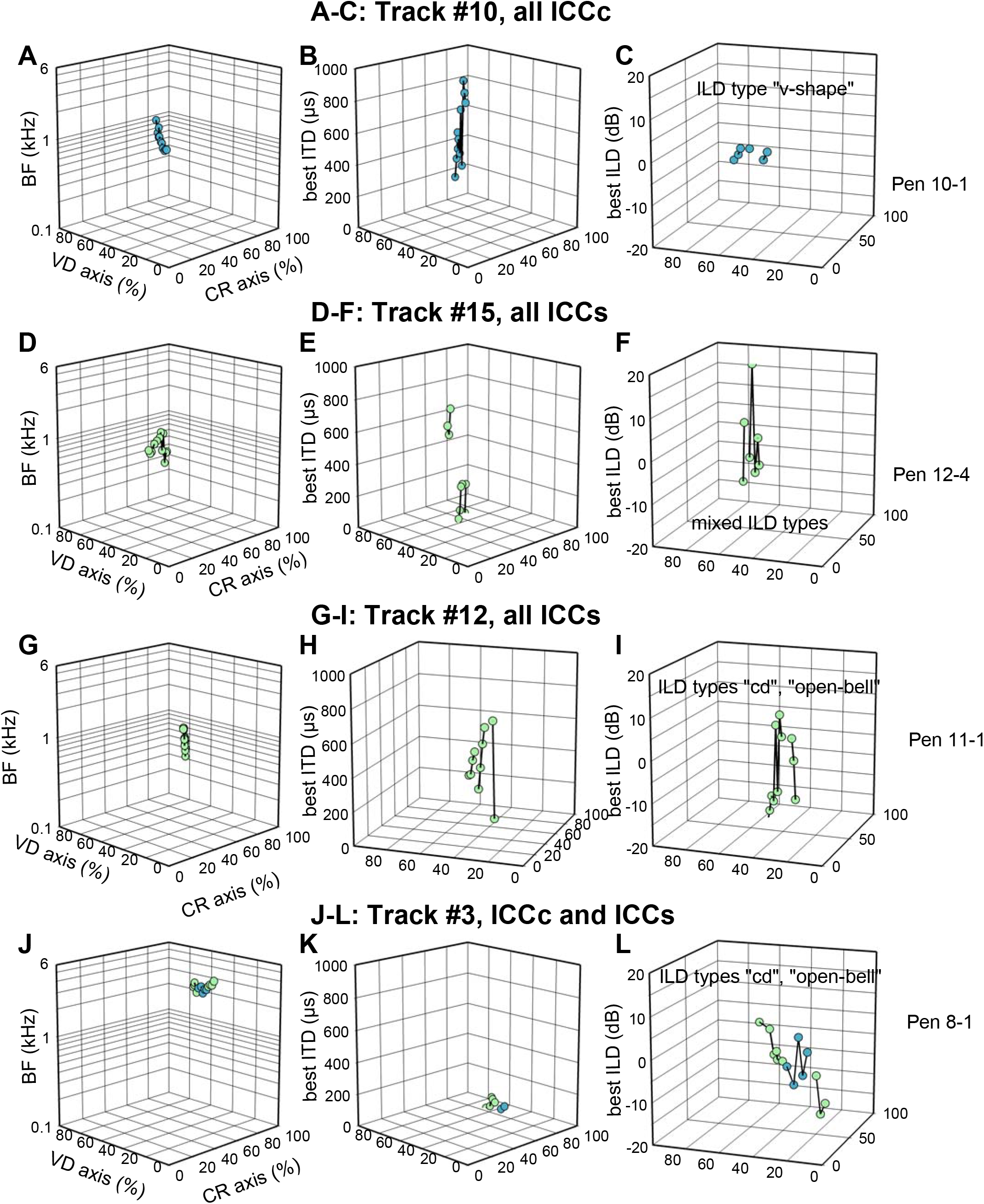
Examples of topographies along individual electrode tracks. Data for 4 different tracks are shown, organized in rows (A-C, D-F, G-I, J-L) and numbered as previously published (Aralla et al., 2020); supplementary material). The 3 columns of graphs illustrate the recording sequences for best BF (A, D, G, J), best ITD (B, E, H, K) and best ILD (C, F, I, L). Each 3D graph plots the physiological parameter along the vertical axis, and relative electrode position along the IC’s ventro-dorsal and caudo-rostral axis on the other two axes. Symbol colour indicates whether the recording site was histologically located in ICCc (blue) or ICCs (green) and successive recording sites are joined by lines.

To probe for any topographies in the ILD responses, we carefully examined plots of best ILD as a function of the 3 anatomical axes, separately for the different ILD response types, and separately for LF and HF units. This revealed only one subpopulation with evidence for a systematic topography of best ILD. HF units of the “contra-dominated” type of ILD selectivity showed values of best ILD that corresponded to ipsilateral auditory space ventrally in the ICC, and this systematically shifted to contralateral space dorsally (Fig. 10; n = 21, rho = 0.58, p = 0.006). Most of these units were located in the ICCs and they alone robustly showed that trend of best ILD (n = 13, rho = 0.76, p = 0.003). An example for an individual electrode track is illustrated in Fig. 9L; note also the example in Fig. 9I, where a track with only LF units showed no ILD topography. Within those HF units of the “contra-dominated” type of ILD selectivity, there was no relation to their best ITD; indeed, less than half were ITD-selective at all. Note also that this topography of best ILD covered both ipsi- and contralateral auditory space, and was only found along the ventro-dorsal anatomical axis of ICC, and there, ran opposite in direction to the azimuthal representation defined by the best ITD of the other unit types described above.

**Figure 10:**
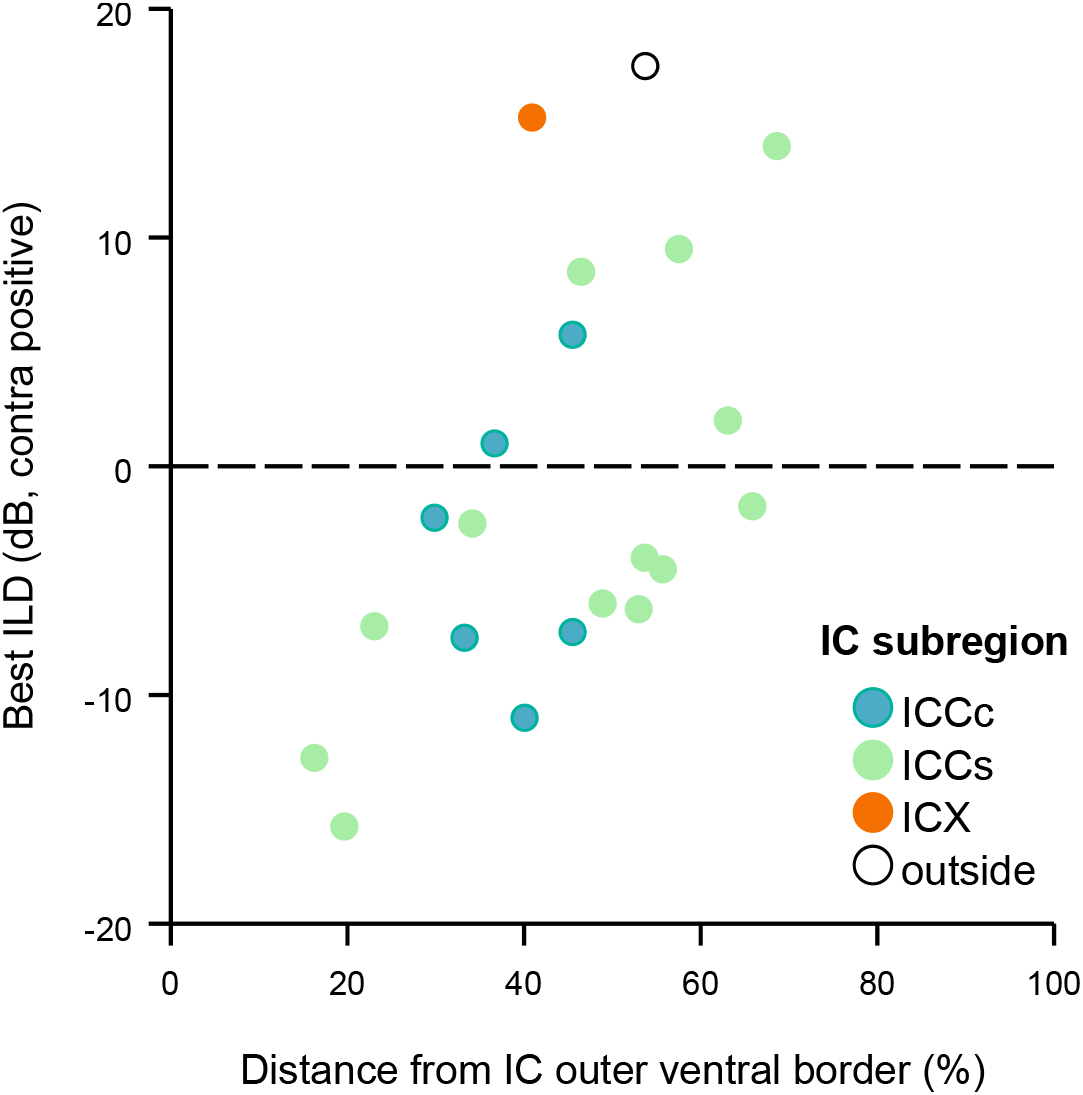
Topography of best interaural level difference. Data are only shown for one selected ILD response type, termed “contra-dominated”. Best ILD is plotted as a function of the recording site’s position along the ventro-dorsal axis. The dotted line indicates zero ILD for reference. Note that the majority of units was located in ICCs (n = 13, shown in green). ICCs units alone, as well as the pooled population, showed a significant correlation of best ILD and ventro-dorsal position, supporting a systematic topography.

In summary, frequency selective units were clearly tonotopically organized, with BFs increasing from rostro-dorsal to caudo-ventral positions in the ICC. Among LF units of both ICC core and shell, there was a clear topography of best ITD, representing the contralateral half of the auditory azimuth, from rostro-dorsal (frontal auditory space) to caudo-ventral. A topography for best ILD was only found in the shell of ICC, and only within HF units of the ILD response type “contra-dominated”. However, this ILD topography was not concurrent with any ITD representation. Finally, due to the paucity of recording sites within ICX, it was not possible to assess any topographies in this subregion.

## Discussion

The core question of the present study was whether the histologically defined subregions of chicken IC match functionally distinct neurone populations based on selectivity for the binaural localization cues ITD and ILD. We found characteristic differences between the core and the shell of the ICC that are graphically summarized in Fig. 11. However, ICC core and shell were not sharply distinguishable, neither anatomically nor physiologically. Below, we will first discuss whether characteristic IC subregions are shared between birds, and thus how well different published characterizations agree. We then ask whether the hierarchical processing of the sound localization cues ITD and ILD in the different IC subregions, as shown in beautiful clarity for the barn owl, is recognizable in less-specialized birds. Finally, we relate IC processing to higher-order multisensory integration and behaviour.

**Figure 11:**
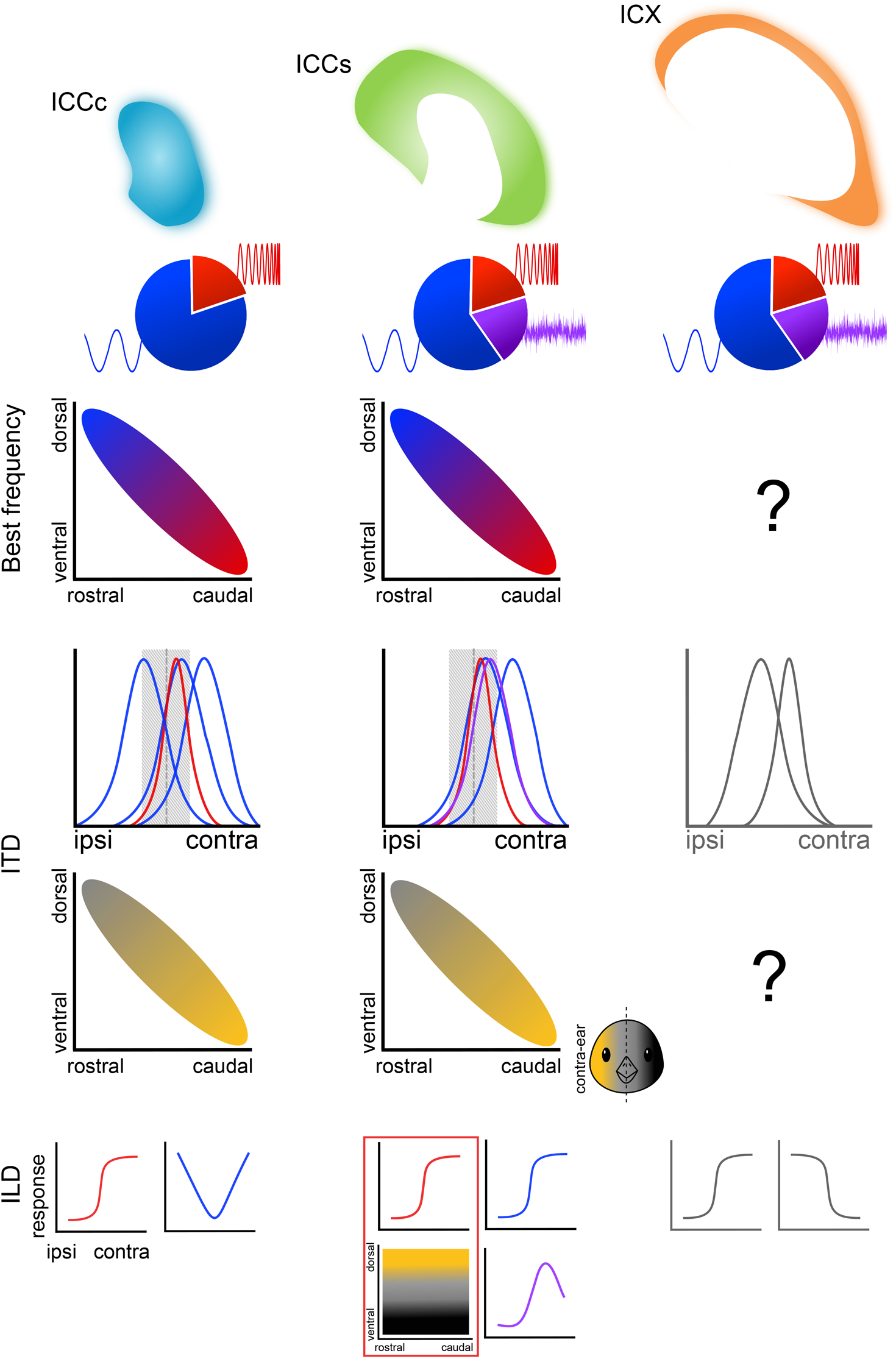
Schematic summary of the distribution of unit response types and topographic representations in the chicken IC. The three subregions evaluated here, ICCc, ICCs and ICX are separated into three columns of graphs. From top to bottom are shown: Typical cross-sectional outlines of all subregions, colour-coded as in earlier figures. Pie charts show the relative proportions of low-(blue) and high-frequency (red) tuned units, and not frequency selective units (purple). The same colour code is used to illustrate the tonotopic gradients along two dimensions, found in ICCc and ICCs. Next, the typical distribution of best ITDs is illustrated by example curves for single LF (blue) and HF (red) units; for reference, gray shading illustrates the approximate natural ITD range heard by the chicken. Below that, the topographic gradients found for best ITD are illustrated which should correspond, in free field, to a map of contralateral auditory space, indicated by the yellow-to-gray shading. The bottom row of graphs illustrates the most common ILD response types of each subregion, represented by schematic single-unit selectivities, and colour-coded to indicate low-(blue) and high-frequency (red) tuned units, and not frequency selective units (purple). Finally, the topographic gradient of best ILD found for units of the “contra-dominated” response type is schematically illustrated using the same colour code as for the best-ITD topographies. Note that this topography is predicted to cover the full ipsi-to-contralateral space in free field.

### No agreed anatomical definition of the subregions in avian auditory midbrain

Our histological analysis identified three main subregions in the chicken IC (Fig. 1 and 2): a core (ICCc) and a shell (ICCs) of the ICC, and the external nucleus (ICX), compatible with the anatomical subdivisions previously established in the chicken (Niederleitner & Luksch, 2012) and in the barn owl (Knudsen, 1983; Wagner et al., 2003). Note that we did not include the superficial ICS in our analysis, which was not expected to be labelled by our routine antibodies (Niederleitner & Luksch, 2012). The use of complementary antibodies was crucial for identifying the different subregions in the IC. As already established in chickens by Niederleitner and Luksch (2012), and confirmed here, the border between ICX and ICCs is distinctly marked by the pair of anti-CKII and anti-PV antibodies. Ideally, it is still desirable to have an additional immunolabel that better highlights the border between the core and the shell of ICC. For this, anti-calretinin was previously found to be suitable in the chicken (Niederleitner & Luksch, 2012; Puelles et al., 1994). However, in our hands, immunoreactions with anti-calretinin could not provide this distinction with sufficient clarity.

The 3-dimensional quantification generated in this study added new information on the relative sizes and spatial positions of the three subregions. Typically, the core was located centrally, in the ventrolateral region of IC, and had an oval disk shape, flattened ventro-dorsally. The shell wrapped almost uniformly around the core, but thickening rostrally, while ICX enclosed the shell with a dorsal thin layer and with lateral and medial wing-shaped regions.

However, the definition and nomenclature of avian IC subregions is not unanimous between studies and may partly depend on the features evaluated. For example, a more varied and different combination of immunolabels led Puelles et al. (1994) to define more, and different, subregions in the chicken IC than Niederleitner and Luksch (2012), whose protocol we followed here. This may also be the reason why neither we nor Niederleitner and Luksch (2012) could distinguish between medial and lateral shell of the ICC, which have been defined physiologically in the barn owl (Adolphs, 1993a), but were also suggested to be distinguishable by chemoarchitectonic criteria in the chicken (Puelles et al., 1994). We hypothesise that our definition of ICCs includes what Puelles et al. (2019; 1994) suggest are three separate regions, i.e. their central shell (CSh), central caudomedial part (CCM) and paracentral nucleus (PCe). There also remains a discrepancy regarding the medial extent of ICX, which typically defined the outer medial border of the IC in our samples, a position clearly taken by the CCM in Puelles et al. (1994; 2019). This strongly suggests different definitions of ICX. Similar discrepancies between different studies have been pointed out for the definition of IC subregions in the barn owl (Wagner et al., 2003). ln the zebra finch, Logerot et al. (2011) appeared to arrive at a similar histological definition of ICCc and ICCs to Niederleitner et al. (2012) in the chicken, although they used a different nomenclature and did not consider areas beyond ICC.

Beside the histological identification, the tracing of brainstem projections to the IC has also been used to define the ICC and different subregions within it. In most birds tested, two distinct pathways originate from NL and NA and terminate in separate ICC target zones. In the pigeon and the barn owl, the projections from NL target a rostro-medial core of the ICC, while those from NA distribute partly around that and more caudally, defining a kind of shell (Leibler, 1976; Takahashi & Konishi, 1988; Wild, 1995). In the chicken, three distinct non-overlapping regions have been defined: a small central “NL-recipient” region within a large “NA-recipient” region, and a dorsal “intermediate” region (Conlee & Parks, 1986; Wang & Karten, 2010). The latter was defined by inputs from a regio intermedia (RI) in the brainstem, adjacent to NL, NA, and NM at their caudal level, as well as from scattered neurones of the VIIIth nerve (Wang & Karten, 2010).

Interestingly, in our samples, the shell of ICC clearly presented different degrees of the labelling intensity of cell bodies and somata, fading toward the core with anti-CKII, while simultaneously becoming darker with anti-PV. Puelles et al. (1994) also observed a very similar labelling gradient using several other markers. This strongly suggests heterogeneity of the shell’s cellular composition. If the histologically defined ICCs is indeed the target zone of NA, heterogeneous inputs to the ICCs might well contribute to this heterogeneity; based on their morphology and physiological responses, NA has at least four or five different cell types (reviewed by (Bloom et al., 2014; MacLeod & Carr, 2007).

In summary, the above discussion highlights the fact that there is currently only limited agreement on the definition of IC subregions in birds, probably mainly due to different techniques and criteria employed. Importantly, the three types of definitions, based on histology, connectivity, or on physiological characteristics, have rarely been brought together in the same study. Therefore, it often remains unclear whether distinct projection target zones or zones of physiologically distinct responses match the subregions defined by immunolabelling and cytoarchitecture. In two different studies on the zebra finch, the same lab characterized the MLd (equivalent to the ICC by their definition) by immunolabelling (Logerot et al., 2011) and also determined brainstem projection zones (Krützfeldt et al., 2010). A comparison suggested that their histologically defined inner and outer MLd bore no relation to the brainstem projections from NL and NA, which, however, did not clearly segregate in the zebra finch.

### Representation of binaural disparities and their presumed hierarchical processing within the avian ICC

In the barn owl, the core of the ICC receives ITD-selective inputs from contralateral brainstem nuclei: the nucleus laminaris (NL; Takahashi & Konishi, 1988) and presumably from the anterior part of the dorsal lateral lemniscus (LLDa; as suggested for the pigeon; Leibler, 1976). The entire spectrum of the owl’s hearing range is represented in the ICCc, with sharp frequency tuning of the individual neurones and a dorso-ventral tonotopic organization. ITD sensitivity still shows phase ambiguity, and systematically represents ipsilateral azimuthal space along an axis perpendicular to each isofrequency band (Takahashi et al., 1989; Wagner et al., 2007; Wagner et al., 2002). However, ICCc neurones typically do not show any true ILD sensitivity, that is, if their discharge does modulate with ILD, it could be explained by linear summation of their bilaterally excitatory inputs (Wagner et al., 2002).

In a nearly complementary fashion, the medial shell of ICC receives projections from contralateral brainstem nuclei involved in sound-level coding: the nucleus angularis, superior olive, and the posterior part of the dorsal lateral lemniscus (LLDp; Adolphs, 1993a; Takahashi & Konishi, 1988). ICCms neurones are sharply frequency tuned, tonotopically organized, and sensitive to ILD but not ITD (Adolphs, 1993a).

At the level of the lateral shell of the ICC in the barn owl, the two independent ascending pathways processing ITD and ILD then converge for the first time, such that the ILD-sensitive inputs from the contralateral posterior part of the dorsal lateral lemniscus (LLDp) are combined with the ascending ITD-selective information from the contralateral ICCc (Takahashi & Konishi, 1988; Takahashi et al., 1989). Similar to ICCc, the lateral shell of ICC presents frequency tuning curves with a narrow-to-intermediate bandwidth and a dorso-ventral tonotopy. However, unlike in ICCc, frequencies below 2.5 kHz are only sparsely represented (Wagner et al., 2007). Furthermore, the topographic map of ITD in the lateral shell now represents contralateral space (Takahashi et al., 1989; Wagner et al., 2007). ILD selectivity is also present in ICCls, with sigmoidal or open-bell-like curves, selective for ILDs louder in the contralateral ear (Adolphs, 1993b). A systematic topography of ILD representation has not been reported.

At the apex of IC processing in the barn owl, the external nucleus (ICX), neurones are much more broadly frequency-tuned, suggesting convergent inputs by a range of narrowly tuned sources (Knudsen, 1984; Knudsen & Konishi, 1978; Wagner et al., 2007). However, a basic tonotopic order is maintained that runs congruent with the topographic map of azimuth, such that frontal auditory space is primarily represented by neurones sensitive to high frequencies and lateral space by neurones with progressively lower-frequency selectivity (Cazettes et al., 2014; Pena et al., 2019). If stimulated with broadband noise, the ITD tuning curves show a dominant main peak and strong side-peak suppression (Fujita & Konishi, 1991; Peña & Konishi, 2000; Takahashi & Konishi, 1986). Furthermore, ICX neurones are selective for a narrow range of ILDs close to zero, such that the ILD-tuning curves are typically bell-shaped (Takahashi et al., 1984).

A number of features in our chicken data were broadly consistent with these classic barn owl data. Units in the ICCc were always frequency selective, their BF distribution tended to be low-frequency biased, and the dominant tonotopic axis also ran ventro-dorsally. Units in the ICCs tended to be more sensitive to high frequencies, and all units with broadband responses were found there. Furthermore, all the rare units with “open-bell-shaped” ILD selectivity were located in the outer shell region. This is in line with the physiology of the lateral ICCs in the barn owl (note that within the ICCs, our recordings almost exclusively originated from the lateral half). However, the chicken data also differed in three important aspects from the classic barn owl data.

First, we found almost no ITD-selective units with best ITDs in the ipsilateral hemifield, as described for the barn owl’s ICCc, and also expected for the chicken. As discussed above, after tract tracing in the chicken, an “NL-recipient zone”, receiving projections from the contralateral brainstem NL, was described (Wang & Karten, 2010). Since the best-ITD topography in NL corresponds to the contralateral acoustic hemisphere in chicken (Köppl & Carr, 2008), at the level of IC this projection was expected to result in best-ITD responses matching the ipsilateral hemisphere. The unexpected paucity of ipsilateral best ITDs could indicate that we did not adequately sample the IC’s NL-recipient zone – which in turn would suggest that Wang and Karten’s (2010) NL-recipient zone does not correspond to our definition of ICCc (based on antibody labelling). However, we note that our findings in chicken are consistent with Lewald’s (1990) large sample of neurones, penetrating the IC in a grid-like fashion in awake pigeon. In the pigeon, too, most IC units were found to respond to contralateral or frontal free-field stimuli, and only a small minority of 6% responded to ipsilateral stimuli (Lewald, 1988). For a subset that was tested with dichotic stimulation, this matched their ITD selectivity (Lewald, 1990). Furthermore, a topographic representation of contralateral auditory space was suggested for the pigeon along the IC’s caudo-rostral axis (Lewald, 1990), which matches our findings for best-ITD representation in the chicken (Fig. 8).

Second, in the chicken, the (lateral) shell of ICC did not show increased proportions of units that were selective for both ITD and ILD. In fact, the physiology of ICC core vs. shell was not distinctly different with respect to the parameters tested here (Fig. 11). Both regions even shared the same principal topographic axes for BF and best ITD. This highlights once more that mismatches exist between different definitions of IC subregions, based on either antibody labelling, tract tracing, or physiology. The present study is the first non-owl work that attempted to match physiology and anatomical subregions in the avian IC.

Third, we found no evidence for any coherent spatial representation that was based on neural selectivities for both ITD and ILD. The only systematic map of best ILD in the chicken did not match or extend the ITD representation when both are translated to spatial coordinates in the free field. Furthermore, the rare recordings located in the chicken’s ICX did not consistently show broadband responses or enhanced selectivity for spatial cues.

### The coding of auditory space and multimodal integration beyond IC

The barn owl’s auditory space map, with azimuthal coordinates based on ITD-selectivity and elevational coordinates on ILD-selectivity, is a unique specialisation associated with outer-ear asymmetry. However, even in the barn owl, this distinction is not absolute (Kettler et al., 2017). It is widely assumed that the barn owl’s specialised case was derived from a plesiomorphic pattern in symmetrically-eared birds, where, according to the duplex theory (Rayleigh, 1907), ITD-selectivity from predominantly low-BF neurones and ILD-selectivity from predominantly high-BF neurones together synthesise a map of auditory azimuth. This assumption is supported by data from symmetrically-eared owl species in which ICX neurones map only azimuth and show no selectivity for auditory elevation. Otherwise, the IC subregions’ selectivities for ITD and ILD are similar to those in the barn owl, which suggests the same hierarchical, step-wise integration of ITD- and ILD-selectivities, culminating in an auditory azimuth map in ICX (Volman & Konishi, 1989, 1990).

Both in our chicken data and in pigeon (Lewald, 1990), a similar processing hierarchy within IC was not evident. On the basis of physiology in pigeon and chicken, no clearly different neurone populations could be identified that would correspond to the owls’ core and shell of ICC, or its ICX. In our chicken data, adding histological identification of these subregions did prove that at least ICCc and lateral ICCs were well sampled, but distinct physiologies did not correlate with that. In both pigeon and chicken, the physiology of IC neurones suggested the coding of the spatial cues ITD and ILD according to the duplex theory: ITD selectivity was only (pigeon) or predominantly (chicken) found in low-BF units, whereas ILD selectivity was only (pigeon) or predominantly (chicken) found in high-BF units (Aralla et al., 2020; Lewald, 1988, 1990). Furthermore, low-BF units represented ITD topographically in both species (Lewald, 1990); this study), and high-BF units formed an ILD topography in the chicken. However, although both topographies coexist in the chicken’s lateral ICCs, they do not correlate and do not form a uniform topography of azimuth, a unity that would be expected in the duplex theory. It may be argued that by not adequately sampling the ICX, we missed the crucial integrative step. However, in the barn owl, and probably also symmetrically eared owls, the integration of ITD and ILD selectivities is a multi-step process that is believed to begin in the lateral ICCs (although the definitions of lateral ICCs vs. ICX may differ between studies (Adolphs, 1993a, 1993b; Mazer, 1998; Volman & Konishi, 1990).

In summary, hierarchical, step-wise integration of the localisation cues ITD and ILD, taking place in different subregions of the IC and culminating in an auditory space map, has not yet been demonstrated in any non-owl species. In the barn owl, the two-dimensional auditory space map in ICX is forwarded by a direct projection to the optic tectum, and merged with an analogous retinal projection, to create a bimodal topography of neurones with restricted visual and auditory receptive fields that are largely in register (Knudsen, 1982). Interestingly, a similar direct projection from ICX to optic tectum hardly exists in the chicken. Instead, the predominant projection is indirect, via an external portion of the formatio reticularis lateralis (FRLx; Niederleitner et al., 2017). Physiologically, very broadly confined auditory receptive fields of two basic types - annular-shaped and circular - predominated in the FRLx and optic tectum, respectively (Maldarelli, Firzlaff, Kettler, et al., 2022). Both types of receptive fields centred around the lateral visual axis of the chicken and differed primarily in size, but not in azimuthal position (Maldarelli, Firzlaff, Kettler, et al., 2022). In FLRx, receptive-field size was also topographically represented (Maldarelli, Firzlaff, Kettler, et al., 2022). However, the auditory receptive fields were all very much larger than the previously recorded visual receptive fields in the optic tectum (Verhaal & Luksch, 2013). Selectivity for ITD was sharper, and the range of best ITDs smaller, than we found in IC, suggesting further sharpening in FRLx and optic tectum. Furthermore, best ITD clearly correlated with receptive-field size, which suggested that ITD selectivity is a major determinant of the auditory receptive field of those neurones (Maldarelli, Firzlaff, Kettler, et al., 2022). Selectivities for ILD were less diverse (only of the type “contra-dominated”) than we found in IC, and not clearly improved. Although a correlation also existed between best ILD and receptive-field size, any contribution to shaping the spatial receptive field remained unclear, questioning neural coding according to the duplex theory.

### Relation to behaviour

The midbrain IC is a major hub of auditory processing and feeds forward into two basic processing streams (Knudsen, 2018). One stream goes to the thalamus and forebrain, ultimately resulting in conscious perception and behavioural decisions. Another stream goes to the midbrain optic tectum (aka superior colliculus) for multimodal integration and ultimately drives fast, reflexive orienting responses to novel stimuli in the environment (Pena & Gutfreund, 2014). Owls show fast, acoustically driven orienting responses with beautiful precision (Knudsen et al., 1979), bringing novel stimuli into the zone of stereoscopic depth vision of their frontally directed eyes (Harmening & Wagner, 2011) and into their frontal acoustic field of most precise neural representation (Pena et al., 2019). Most other birds, like the chicken, do not show such precise orienting responses, indeed, probably do not need them, that is, there was no selective pressure for it. Most birds are diurnal and predominantly visually oriented, and their laterally directed eyes provide a near-panoramic view of the environment (Martin, 2017). As argued before (Maldarelli, Firzlaff, Kettler, et al., 2022; Niederleitner et al., 2017), the auditory contribution to fast orientation towards novel stimuli is probably minor and the midbrain circuits of the chicken reflect that.

Note that this does not mean that chickens do not pay attention to auditory stimuli. In fact, chickens are surprisingly precise in conditioned behavioural sound localization tests, although it can be difficult to train them and achieve proper stimulus control (Krumm et al., 2022; Maldarelli, Firzlaff, & Luksch, 2022). This further supports the notion that, for the chicken, the auditory sense is not a dominant driver of fast, reflexive orienting responses, but is more important for conscious behaviours driven by context and motivation.

## Materials and Methods

### Experimental Animals

Seven chickens (*Gallus gallus*), of both sexes and aged between 21 and 36 days post hatching (P) were used. Five animals were hybrids of male red junglefowl (*Gallus gallus bankiva*) and female Zwerg-Welsumer (a variety of *Gallus gallus domesticus*) from a colony at the animal facility of the University of Oldenburg. The remaining two chickens were of the commercial breed Selected White Leghorn (Gallus gallus domesticus) that hatched from pathogen-free eggs. All animals hatched and were raised at the same animal facility and were therefore exposed to the same environment. We did not observe any differences in auditory physiology between the two breeds (Aralla et al., 2020).

### Electrolytic lesions and histology

During the electrophysiological experiments (Aralla et al., 2020), a subset of recording sites was marked with electrolytic lesions for the histological reconstruction of the electrode tracks. Typically, two lesions were placed along selected electrode tracks after the recordings of neural activity, one at a location near the start and one near the end of auditory responses. The lesions were generated by disconnecting the amplifier so that positive DC current (10 µA) could pass through the electrode for 20 seconds via a lesion maker (Ugo Basile SRL, Biological Research Apparatus, Varese, Italy). In a given brain, between one and four electrode tracks were thus lesion-marked. Once the experiment was concluded, the animal was euthanized with an overdose of sodium pentobarbital (“Narkodorm”, CP-Pharma GmbH; 1 ml/kg), then perfused transcardially. The perfusion started with 100 ml of phosphate-buffered saline (PBS) flushing out the blood, followed by 0.2 ml heparin in 150 ml PBS, and finally 250 ml of 4% paraformaldehyde in PBS to fix the tissues. After perfusion, the brain was extracted from the skull and cut along the midline into two hemispheres. The cerebellum was then dissected away and the forebrain partially removed by a vertical cut orthogonal to the midline, defining the future plane of section. Importantly, the ventro-dorsal orientation of the section plane was the same as in the brain atlas by Puelles et al. (2019) and also aligned with the vertical electrode axis in our stereotaxic setup. The two brain hemispheres were then immersed in 30% sucrose in PBS for 48 h for cryoprotection and subsequently frozen in crushed dry ice and stored at -80°C. Frozen 30-µm thick cross sections were obtained using a cryostat (Leica CM 1950; Leica Biosystems, Wetzlar, Germany) and individually collected in PBS in multiwell plates. Sections containing IC, plus a few sections before and after, were selected using a stereomicroscope and were assigned to different labelling protocols.

Alternating sections were typically selected for Nissl or immunohistochemical labelling. Usually, one series was immediately mounted on gelatine-coated slides and Nissl-stained with cresyl violet to visualize the whole IC and the electrolytic lesions. Immunolabeling was performed on free-floating sections (all steps with gentle agitation), with different primary antibodies, as detailed in Table 3. The choice of antibodies and their concentrations was guided by published work (Niederleitner & Luksch, 2012; Puelles et al., 1994; Wagner et al., 2003) and previous labelling tests carried out in our laboratory. We aimed at employing antibodies that resulted in complementary labelling of IC subregions of interest, to facilitate identification of the subregions’ borders. To this end, we mainly used anti-parvalbumin (PV; monoclonal mouse IgG1, P3088, Sigma-Aldrich), diluted 1:16000, in combination with anti-Ca^2+^/calmodulin-dependent protein kinase II α (CKII; monoclonal mouse IgG1, 05-532, clone 6G9, EMD Millipore Corporation, Temecula, USA), diluted 1:4000, in PBS with 3% bovine serum albumin and 0.2% Triton X-100 added (Table 3). One brain was stained with anti-calretinin (CR; polyclonal rabbit, C7479, Sigma-Aldrich, USA), diluted 1:2000 and counter-stained with Nissl staining. Incubation with primary antibodies was for at least 24 hours, at 8°C. After several washes in PBS, the sections were incubated with a biotinylated secondary antibody (anti-mouse: horse, BA-2000, Vector Laboratories, Burlingame, USA; anti-rabbit: goat, B8895, Sigma-Aldrich, USA), diluted 1:1000, for at least 2 hours at room temperature. Avidin–biotin–peroxidase complex binding (Vector Labs, 1:500, 1h) was performed next, followed by a nickel-intensified enzymatic conversion of diaminobenzidine (DAB; 0.03%, 100mM NiSO4, 0.003% H_2_O_2_ in a buffer of 175 mM Na-acetate and 10mM imidazole).

**Table 3:** List of the staining procedures and the antibodies used for each sample.

| Animal | Staining | Primary antibodies | Secondary antibodies |
| --- | --- | --- | --- |
| **RAIC6** | Nissl |  |  |
| **RAIC8** | Nissl |  |  |
| **RAIC9** | Nissl, DAB | anti-calretinin (rabbit) 1:2000 | anti-mouse-Biotin (horse) |
| **RAIC10** | Nissl, DAB | anti-CKII (mouse) 1:4000<br>anti-parvalbumin (mouse) 1:16000 | anti-mouse-Biotin (horse)<br>anti-mouse-Biotin (horse) |
| **RAIC11** | Nissl, DAB | anti-CKII (mouse) 1:4000<br>anti-parvalbumin (mouse) 1:16000 | anti-mouse-Biotin (horse)<br>anti-mouse-Biotin (horse) |
| **RAIC12** | Nissl, DAB | anti-CKII (mouse) 1:4000<br>anti-parvalbumin (mouse) 1:16000 | anti-mouse-Biotin (horse)<br>anti-mouse-Biotin (horse) |
| **RAIC16** | Nissl, DAB | anti-CKII (mouse) 1:4000<br>anti-parvalbumin (mouse) 1:16000 | anti-mouse-Biotin (horse)<br>anti-mouse-Biotin (horse) |

### Histological identification of IC subregions and lesions

All sections containing IC and lesions were documented with a slide-scanning microscope (AxioScan.Z1, Carl Zeiss Microscopy GmbH, Jena, Germany) and exported in high resolution to TIFF format, with XY pixel dimensions of 0.441 µm. These images were then loaded into Fiji (“Fiji is just ImageJ”; (Schindelin et al., 2012), and the borders of IC and its subregions (the core (ICCc) and the shell (ICCs) of the central part of IC, and the external nucleus (ICX)) were manually outlined using a tablet and digital pen (Wacom Intuos Pro 430 x 287 x 8mm, Wacom Pro Pen 2). Care was taken to establish nuclear borders with precision, for example by applying overlays of neighbouring sections with a complementary label. Electrolytic lesions were similarly defined and color-coded to ease their identification across sections.

### Three-dimensional quantification of IC and its subregions

The positions and dimensions of IC and its subregions were quantified in three dimensions: medio-lateral, ventro-dorsal and caudo-rostral. First, we obtained distances of the outer nuclear borders to a reference element. In a second step, these border points were converted into normalized coordinates relative to the total extent of the respective structure along each axis. Finally, the normalized border points were used to calibrate the entire nuclear outlines and derive a normalized 3D coordinate space within which the positions of lesions and electrophysiological recording sites could be expressed and compared across brains. More detailed descriptions of the measurements for each axis follow below.

#### Medio-lateral extension

Medio-lateral measurements were obtained only for the total IC in cresyl-violet stained sections, and, in addition, for ICCc, ICCs, and ICX in immunohistochemically labelled ones. Two lines, extending orthogonally from the hemisphere’s midline, were used to define the minimal and maximal distances of a structure’s borders from the midline, with the use of Photoshop (Adobe Photoshop CS6, version 13.0.0.0; example in Fig. 12A). This created a reference grid overlay that could be used later to rotate each image to vertical orientation of the midline. The software Fiji was used to measure the absolute distances to the midline. For each brain, the smallest medio-lateral distance across all sections was then defined as the “null” point, and the largest distance as the maximum, and all measurements were converted to normalized relative coordinates between those extremes.

**Figure 12:**
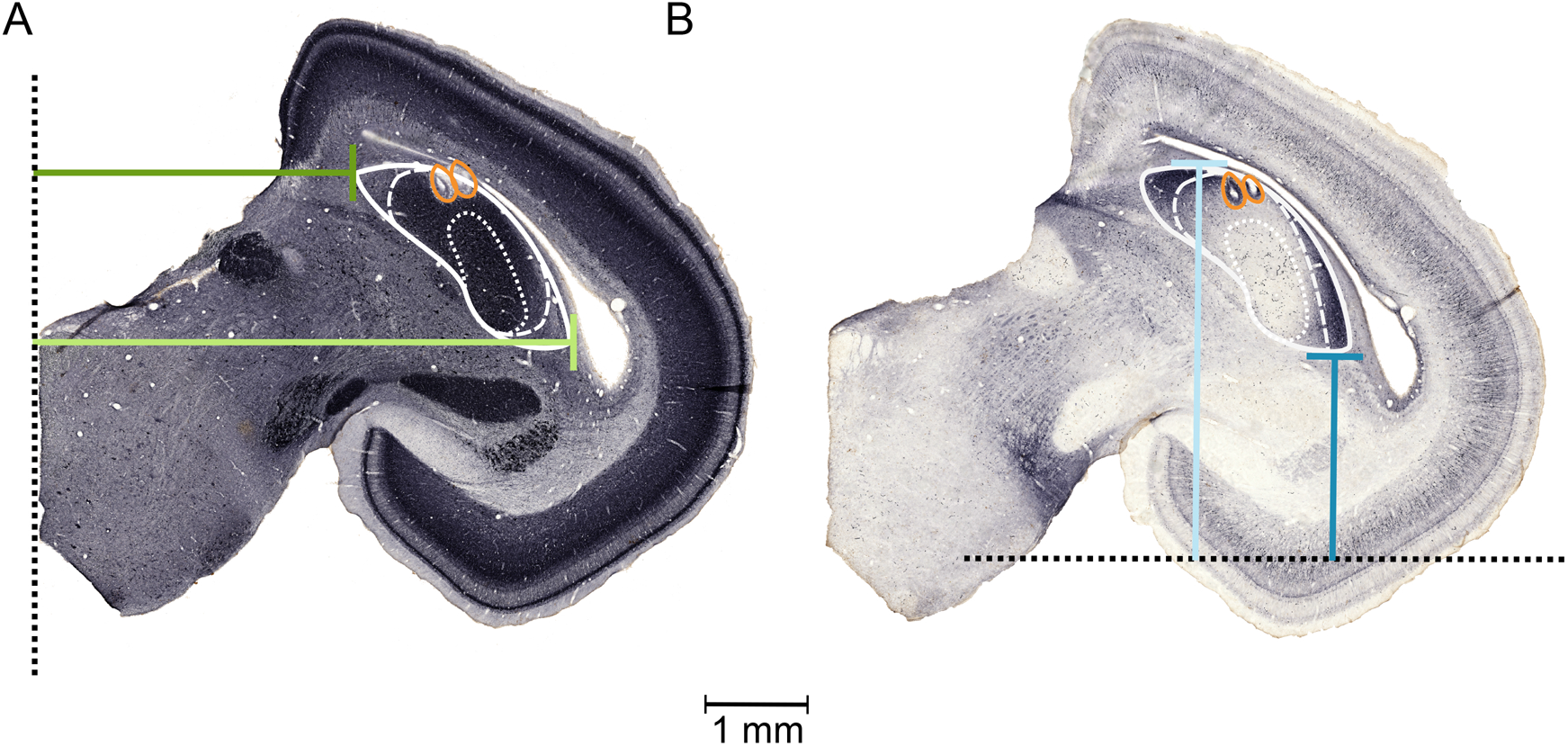
Illustration of the definition of IC’s outer borders. Panels A and B show histological cross-sections of a right midbrain labelled with parvalbumin (A) and CKII (B). The solid white outlines identify ICX, the dashed lines ICCs and the dotted lines ICCc. Orange contours indicate electrolytic lesions. In A, the vertical dotted black line represents the brain’s midline, used as reference for the medio-lateral plane; the minimal (dark green) and maximal (light green) distances to IC’s outer borders (here, represented by ICX) to this reference are also shown. In B, the same is illustrated as dark blue and light blue lines for the ventro-dorsal axis. Here, the horizontal dotted black line represents an arbitrary reference, established after alignment of all relevant sections with the aid of 3D-reconstruction software (see Methods).

#### Ventro-dorsal extension

The borders of IC and its subregions were similarly defined in each section with lines running parallel to the brain’s midline, i.e. the ventro-dorsal axis (example in Fig. 12B). However, we had no absolute ventro-dorsal reference point across sections that would allow their unambiguous alignment. As a solution, a 3D-reconstruction software suitable for the analysis of serially sectioned histological specimens (Voloom V3.0.0, microDimensions Voloom v 3.0.0.) was used for automatic alignment with manual adjustment, and an arbitrary reference line was defined across all aligned sections of a particular brain. From coronal views of the aligned sections, distances to the reference line (dashed line in Fig. 12B) of the ventral and dorsal outer borders were then measured for all structures of interest. Finally, normalization of coordinates along the total ventro-dorsal extent was achieved in the same way as described above for the medio-lateral dimension.

#### Caudo-rostral extension

Caudo-rostral dimensions were given by the number of sections showing IC or its subregions, multiplied by 30 µm (the thickness of each section). Missing sections (if any) that had been lost during histological processing were included in the count, to obtain the correct total size of the brain sample. Normalized coordinates were again calculated relative to most caudal and rostral points, as established for the medio-lateral and ventro-dorsal axes.

### 3D-reconstruction of the subregions and of the electrode tracks

A crucial aim of this study was the three-dimensional reconstruction of the IC and its subregions, to place all brain samples into a common, normalized 3D space. For this final step, we designed a custom script in Matlab (vR2018b; MathWorks, Natick, MA, USA) that loaded all images of a particular brain, rotated them (if necessary) to align the hemispheric midlines across sections, and detected the manually defined nuclear outlines (ICCc, ICCs and ICX). The outlines’ coordinates were then normalized in three dimensions, using the previously determined, normalized outer borders of each structure, in each section, along all three axes (see above), as calibration points. Thus, the subregions’ anatomical borders could then be plotted in a three-dimensional manner. Two early brains that had not been immunolabelled (see Table 3) presented a special case, because the subregions within the IC could not be clearly defined on the basis of the cresyl-violet stain alone. Therefore, the normalized coordinates for ICCc, ICCs and ICX were imported from another (reference) brain.

For reconstruction of the electrode tracks from the electrophysiological experiments, the linear distance between two corresponding lesions in 3D space was calculated. Since the distance travelled by the electrode between two lesions was also known from the readings of the piezoelectric motor during the electrophysiological experiment (Aralla et al. 2020), every individual recording site along the track could thus be assigned normalized coordinates in all three dimensions and allocated to a specific IC subregion. We assumed the centre of the lesions corresponded to the position of the electrodes’ tips when passing current. We made no attempt to recover recording sites from tracks that were not lesion-marked or if both lesions belonging to a particular track could not be recovered histologically. If the track was missing only one lesion, we used one of two options: (1) if some slight damage caused by the electrode entering on the midbrain’s surface could be identified, this was used as the second reference point defining the track, (2) if another track was reconstructed in the same brain, with the same electrode angle, we derived a parallel trajectory that passed though the single lesion.

### Categories of reliability in electrode track reconstruction

Since several recording tracks were typically lesion-marked in a given brain, ambiguities could arise when reconstructing electrode tracks. We therefore defined categories of different reliability of this reconstruction, based on the identification of the lesions and their match to the stereotaxic coordinates of electrode tracks in the corresponding electrophysiological recording session.

Important criteria were (1) whether the total number of expected lesions could be found; (2) whether their relative locations on the three anatomical axes matched the known stereotaxic coordinates from the experiment, and (3), more precisely, whether the medio-lateral and caudo-rostral track angles inferred from a lesion pair matched the known stereotaxic electrode angles. Two categories of reliability were defined: “Unambiguous”, if two reference points could be clearly identified and unambiguously allocated to an electrode track, and ”plausible”, if a small uncertainty remained, e.g., if not all lesions could be recovered in a given brain but the missing one could be plausibly estimated; or if the IC’s subregions were defined by importing the coordinates from another sample (for brains stained only with cresyl violet). Furthermore, even if located along an unambiguously recovered electrode track, we downgraded the location of individual recording sites to “plausible” when it fell into a region of uncertainty where the boundary between two subregions had to be interpolated between physical tissue sections.

## Acknowledgements

We thank Carolin Jüchter, Sina Engler and Habibazaman Monir for help with histology and microscopy. Supported by the Deutsche Forschungsgemeinschaft (DFG), via the Collaborative Research Center (CRC ‘Active Hearing’, project A14) and the Cluster of Excellence 1077 ‘Hearing4all’. Language services were provided by stels-ol.de.

